# Reconstruction of nine thousand years of agriculture-based diet and impact on human genetic diversity in Asia

**DOI:** 10.1101/747709

**Authors:** Srilakshmi M. Raj, Allison Pei, Matthieu Foll, Florencia Schlamp, Laurent Excoffier, Dorian Q. Fuller, Toomas Kivisild, Andrew G. Clark

## Abstract

Domestication of crops and animals during the Holocene epoch played a critical role in shaping human culture, diet and genetic variation. This domestication process took place across a span of time and space, especially in Asia. We hypothesize that domestication of plants and animals around the world must have influenced the human genome differentially among human populations to a far greater degree than has been appreciated previously. The range of domesticated foods that were available in different regions can be expected to have created regionally distinct nutrient intake profiles and deficiencies. To capture this complexity, we used archaeobotanical evidence to construct two models of dietary nutrient composition over a 9000 year time span in Asia: one based on Larson et al. (2014) and measured through composition of 8 nutrients, and another taking into account a wider range of crops, cooking and lifestyle variation, and the dietary variables glycemic index and carbohydrate content. We hypothesize that the subtle dietary shifts through time and space have also influenced current human genetic variation among Asians. We used statistical methods BayeScEnv, BayeScan and Baypass, to examine the impact of our reconstructed long-term dietary habits on genome-wide genetic variation in 29 current-day Asian populations (Figure S1, Figure 1, Figure 2). Our results show that genetic variation in diet-related pathways is correlated with dietary differences among Asian populations. SNPs in five genes, *GHR, LAMA1, SEMA3A, CAST* and *TCF7L2*, involved in the gene ontologies ‘salivary gland morphogenesis’ and ‘negative regulation of type B pancreatic cell apoptotic process’ suggest that metabolism may have been primary targets of selection driven by dietary shifts. These shifts may have influenced biological pathways in ways that have a lasting impact on health. We present a case that archaeobotanical evidence can provide valuable insight for understanding how historical human niche construction might have influenced modern human genetic variation.

## Results

The dietary models were constructed to capture different aspects of long-term nutritional information. They incorporate the following unique features: 1) an estimation of a preagricultural diet base, which we term a ‘hunter-gatherer diet’; 2) proportionate variation of different dietary components; 3) variation in the length of exposure to dietary components; 4) inclusion of different food processing techniques to obtain nutritional estimates; 5) the use of glycemic index and carbohydrate content to measure nutrition, as both highlight the impact of increased carbohydrate consumption through time and may play a role in diabetes risk [1–3].

The first dietary model we constructed examines relative differences in the change of nutrient levels in each of the three ADMIXTURE-defined groups (Figure S1). In the second model, unlike the first, we incorporate information on pre-domestication diet by including a baseline dietary model for our populations. We assume that the content is the same across populations, but that the proportion of this diet in each population for each period of time varies. We assume the same nutritional content in each pre-domestication diet primarily to measure the changes brought by domestication. Each of the 29 populations had an individualized reconstruction in Model 2 (Table S9). For example, the Dai population, an agricultural population residing at the border of China adjacent to Laos, Thailand, Vietnam and Myanmar, had dietary history divided into three time-slices: 9000-7000 YBP, 7000-5500 YBP, 5500-0 YBP. The first time slice is an estimated hunter-gatherer diet. In the second time slice, we incorporate millet porridge and diet diversification. By the final time slice, more common carbohydrate-based crops are apparent in the diet, such as short-grained brown rice (Table S9).

### Southwest Asian populations show higher micro- and macro-nutrient abundance than East and South Asians

Our primary dietary model (see Methods) infers that the Southwest Asian populations (mostly sampled from present-day Pakistan) had a greater increase in carbohydrates, lipids and protein content due to domestication events from 9000 years ago. In contrast, South Asian populations showed the latest and at least these three major dietary changes in their diets, as domestication started later here. The greatest nutrient increases in all populations due to diet changes were carbohydrates, followed by protein, then lipids. Separated into segments of 3000 years, consumption of all three components from the domesticated diet increased over time, following known increased adoption of domestication practices throughout Asia (Figure 2a, Tables S1-8).

**Figure 1.**
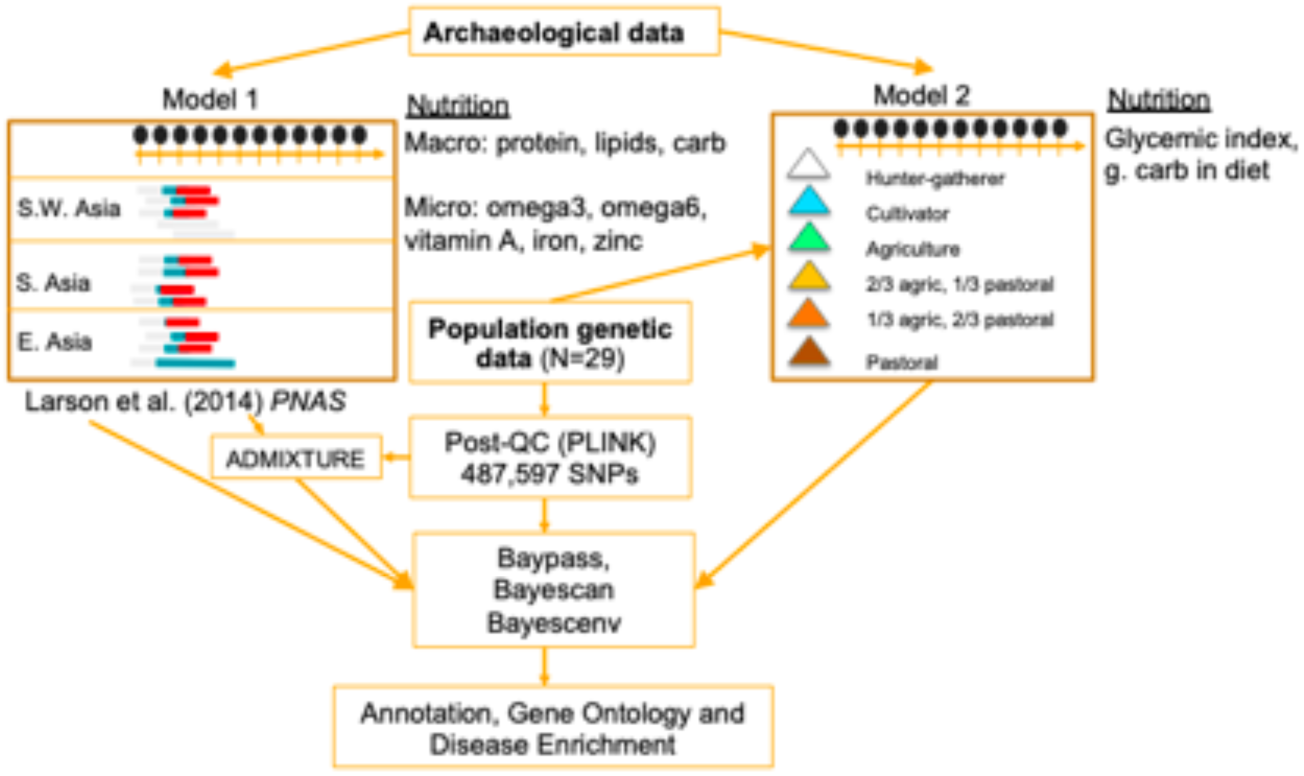
Organization of the study and analysis pipeline. The two figures in column ‘Archaeological data’ refer to the two constructed dietary models, with the boxes beside them referring to the dietary variables from each model.

**Figure 2.**
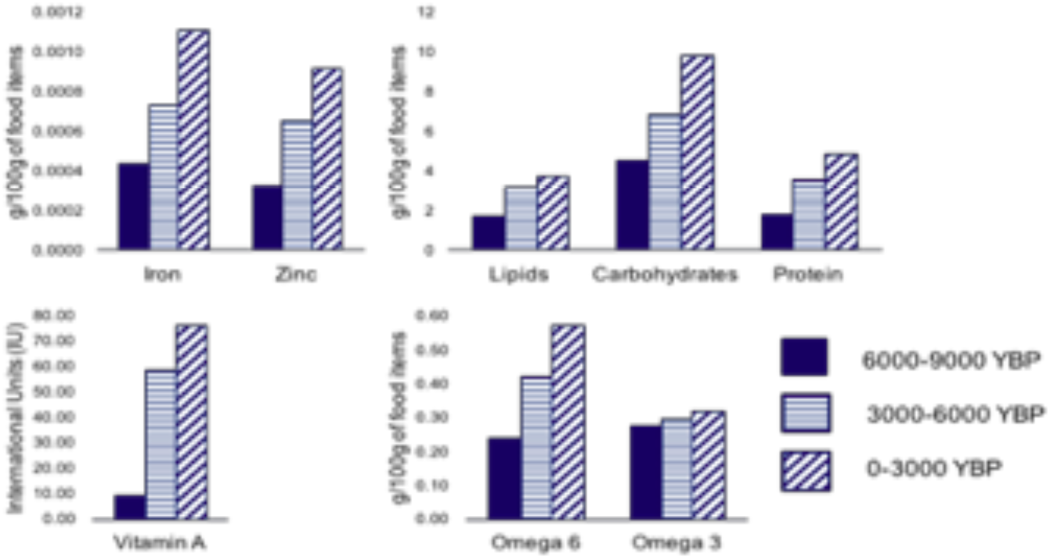
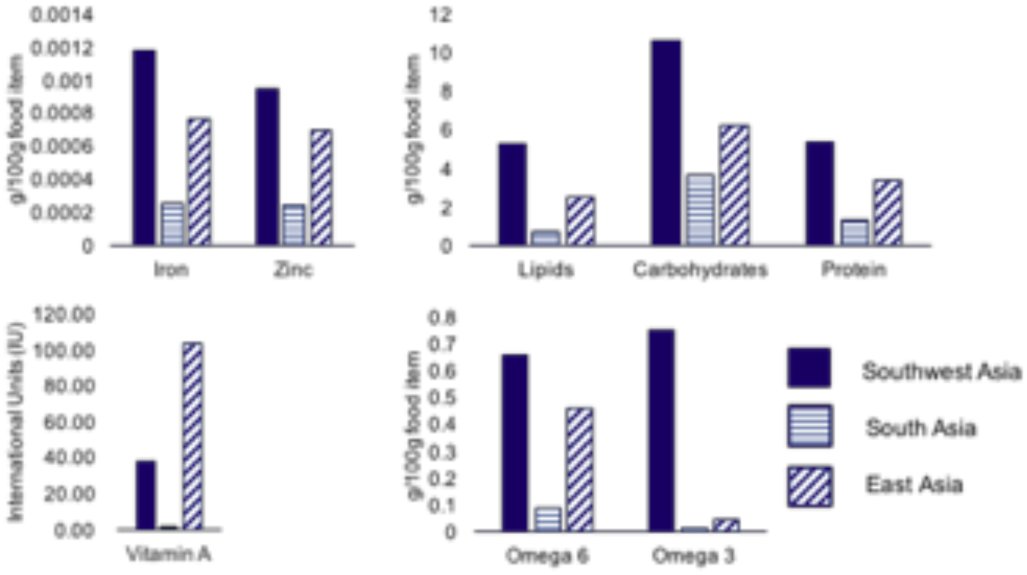
Summary of Diet Model 1. a) Levels of nutrients contributed by domestication, based on Diet Model 1, as a sum total of values across all three population groups. b) Average levels of nutrients across the three time intervals (9000 YBP-present) in each region, based on Diet Model 1. Supplementary Tables 1-8 contain the data used to generate these figures.

Similarly, micronutrients zinc, iron, omega-3 and omega-6 fatty acid content from the diet due to domestication were higher in Southwest Asian populations, followed by East Asia and South Asia populations. The most dramatic differences were observed for omega-3 fatty acids, with Southwest Asia showing higher increases in their diet over the 9000 year duration compared to the other populations. South Asian populations (excluding the Southwest Asian populations, mostly located in present-day Pakistan) showed lower amounts of each of these four nutrients compared to the other two populations (Figure 2b, Tables S1-8).

These results complement evidence suggesting that domestication brought about increased abundance of certain nutrients in the diet, such as carbohydrates, although without a baseline pre-domestication measurement it is difficult to infer from our data alone the magnitude of the increase[4]. Our dietary model does not refute evidence that agriculture also brought about micronutrient deficiencies[5]. Those changes may be modeled with incorporation of caloric content changes, which were outside the scope of this paper. We solely examined changes that occurred in an aspect of agricultural impact on the diet over time. Our finding that micronutrient composition increased over time may reflect the paucity of domestication events at the beginning of the 9000 year window. Furthermore, it is important to note that we did not include a baseline level diet to examine changes due to domestication within each population. It is likely that a forager lifestyle provided a substantial part of the diet during this time. The primary purpose of our constructed dietary model was to compare the nutritional impact of domestication across the populations, through time. The caloric and nutritional content coming from predomestication diet, and species (both native and non-native) that are not included in Larson et al. (2014) are not included in Model 1.

### Modeling intra – and inter-population variability in carbohydrate consumption

In the second model, farming communities (agricultural and cattle herding) had a higher carbohydrate content in their diet due to domestication, across all ancestry groups. However, East Asian agricultural populations had lower levels of carbohydrate content compared to South Asian and Southwest Asian populations. We found substantial geographic and population-level variability in dietary habits and nutritional outcomes across subsistence groups, as we used fine-scale geographical information to construct population diet (Figure 3). Hunter-gatherers’ diet showed lower levels of carbohydrate content, but similar levels of glycemic index to agricultural populations, due to higher presence of fruit in our models. The pastoralist populations showed very similar glycemic index values and carbohydrate content to other populations with both geographic and genetic similarity, i.e. ancestry group 2. We saw a similar inverse relationship between glycemic index and carbohydrate content, across ancestry groups. For example, ancestry groups 2 and 4 show higher carbohydrate content, but lower average glycemic index. Stable agriculture and crop husbandry were responsible for those increases in carbohydrate content (Figure 3, Tables S9-S11).

**Figure 3.**
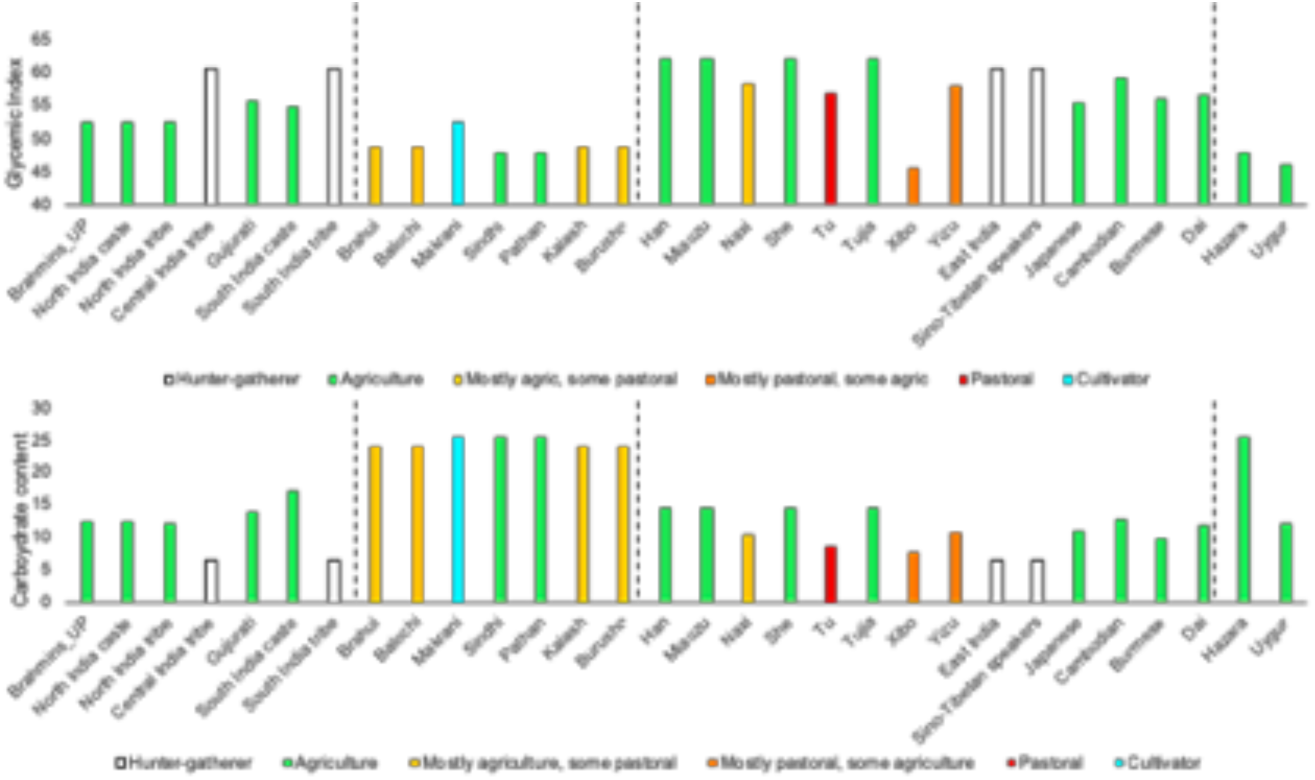
Summary of Diet Model 2. Summary of glycemic index values and carbohydrate content values calculated for each population, based on Diet Model 2. The dotted lines represent population groupings made based on the ADMIXTURE plot (Figure S1), with the first group corresponding to South Asia, second to Southwest Asia, third to East Asia, and fourth an intermediate between Southwest and East Asia (grouped into Southwest Asia). Supplementary Table 10 contains the values for this plot, and Supplementary Table 9 contains the full Diet Model 2.

The carbohydrate content in the second dietary model will differ from the first because the dietary composition in Model 1 referred only to the crops in Larson et al. (2014), and carbohydrate consumption were tailored to specific populations in Model 2, incorporating pre-agricultural diet as well.

### Little overlap between selection tests, but common gene ontology pathways across tests and nutrients

We subsequently analyzed the top 1% of variants from the three Bayesian tests (see STAR Methods) for variants that have been reported to be associated with dietary adaptations, and for overlaps in the results for the three Bayesian tests. Analysis of the top 1% of the variants showed no statistically significant overlap between the three tests (Figure S2). Genes *FADS1/FADS2, OCA2, TYR, LRP2, ADH1B* and *CYP24A1*, previously reported to be under diet-mediated selection, appear in one or more of the top hits for several dietary variables in both dietary models (Figure S3).

All three scans showed similar breakdown of SNPs in each functional class across methods and nutrient variables, except BayeScEnv signals for carbohydrate content (Figure S4). Baypass results had the greatest number of SNPs with deleterious PolyPhen and SIFT scores, but this is not surprising as there were greater numbers of selection results for Baypass compared to BayeScEnv and BayeScan (Figure S5); none of the tests showed enrichment for deleterious or damaging variants.

The number of ontology terms that were in common either among multiple dietary variables or multiple statistical methods for a single dietary variable also varied (Table S12). For example, ‘loop of Henle development’, a gene set involved in kidney development, appeared to be the most strongly enriched term in the BayeScan analysis (binomial FDR q-value: 4.23 × 10^−8^). This ontology term also appeared among the top twenty pathways for vitamin A in both BayeScEnv and Baypass tests (binomial FDR *q*-value: 1.66 × 10^−08^ and 7.07 × 10^−3^, respectively) (Table S12). Retinoic acid, the active form of vitamin A, is critical for most organ development, including the kidney – studies in rats show that even mild vitamin A deprivation can result in kidney underdevelopment [6].

Three other significantly enriched pathways of note include ‘regulation of type B pancreatic cell apoptotic process’, ‘response to food’ and ‘branching involved in salivary gland morphogenesis’. Because GREAT takes entire gene regions into account, we returned to the list of SNPs in the top 1% of results for each of the three tests to identify the individual SNPs located within the genes included in the ontology terms. From these analyses, we found that genes *CAST, TCF7L2, IL6, GHR, LAMA1* and *SEMA3A1* contained SNPs that were located in the top 1% of hits for all three methods, and across multiple dietary variables (Table S12). The genes implicated by ontology terms also play a role in dietary response [7,8][9–11][12][13][14,15][16]. Finally, deletions in the gene *SEMA3A* result in impaired olfactory development, as well as pubertal failure [17], and mutations in the gene can result in pubertal delay [18].

We performed an additional enrichment analysis for selection results in genome-wide association studies. Analysis using the program traseR [19] showed that Baypass results had a greater number and strength of enrichment across disease classes than either BayeScEnv or BayeScan (Figure S6). Most of the significant enrichments were found in anthropometric, metabolic, cardiac, or nutrient-related traits. However, the nutrient-related traits did not always correspond to the specific nutrient variable. For example, ‘Carbs’ (from Model 1) and ‘GI’ showed significant enrichments and larger combined “Odds ratio” for the traits Calcium and Magnesium, respectively, in Baypass results. Body height was the only trait to show significant enrichment for Baypass results across all nutrient variables.

### Allele frequencies of genes among geographically dispersed populations associated with long-term dietary habits are linked to type 2 diabetes risk

Maps of derived allele frequencies for the SNPs associated with the genes and ontology terms in Table S12, Figure 4, Figures S9 and S10 display extensive geographic-region specific stratification. The derived alleles of the vitamin A associated variants in *CAST* and *GHR* genes show higher frequencies in South and West Asia, while the derived and protective[20] allele of *TCF7L2* variant rs10885409 shows the opposite pattern (Figure 4). There is a possibility that the derived alleles have a selective advantage in the geographic regions with higher allele frequencies. Similarly, the *LAMA1* and *SEMA3A* variants show opposite patterns, with the *LAMA1* derived allele showing low frequencies in East Asia followed by West and South Asia, and the *SEMA3A* variant showing the opposite for each of lipids, omega-6, protein and zinc (Figures S9 and S10). Multiple dietary factors (lipids, omega-6, protein and zinc) showing the same trends may suggest a common underlying dietary component exerting a selective pressure on these populations. The opposing directionality of the selective pressure based on the opposite derived allele frequency patterns for *LAMA1* and *SEMA3A* variants suggests that the same dietary components may be beneficial in different dietary contexts.

**Figure 4.**
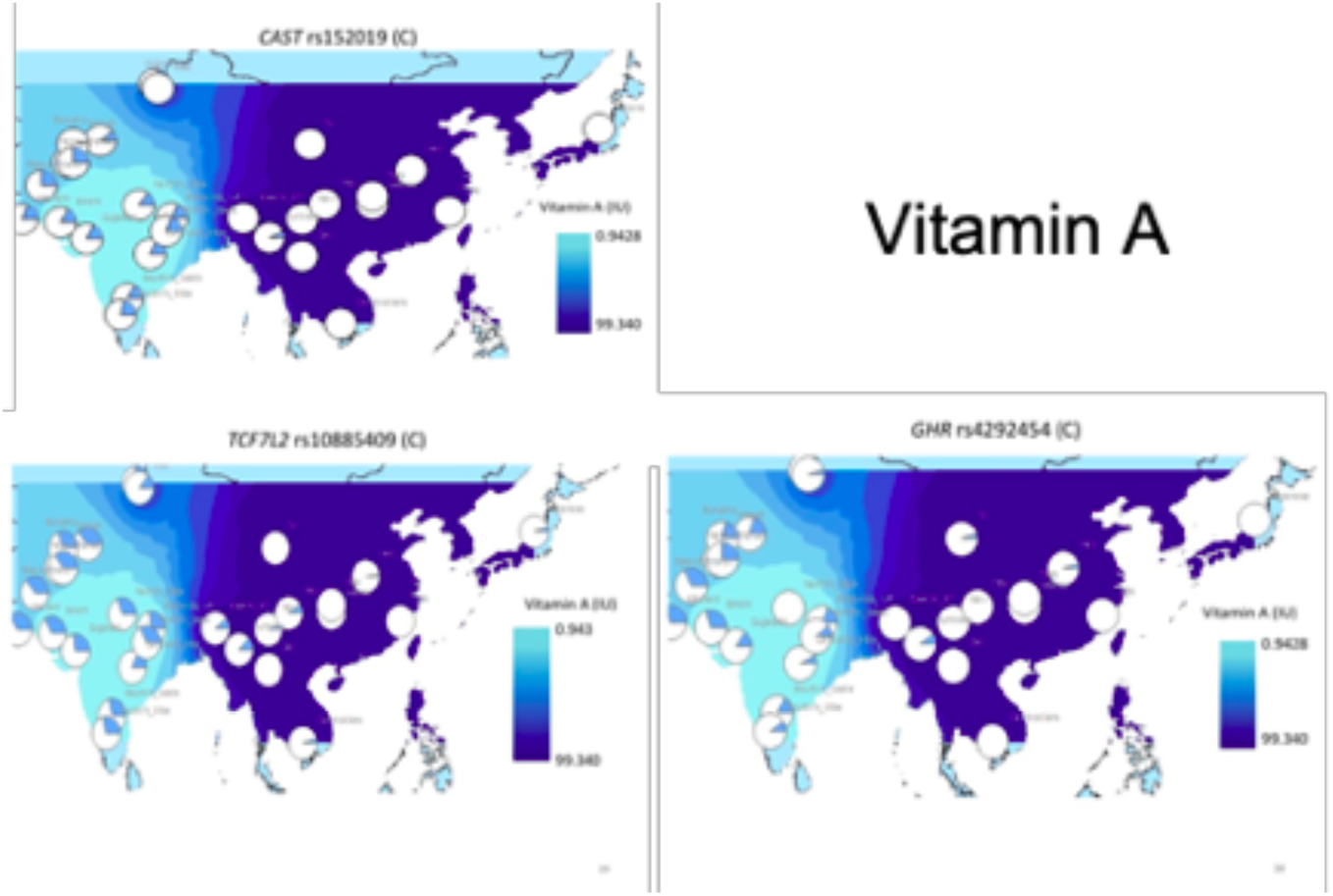
Derived allele frequency and geographic patterns of dietary vitamin A. The pie charts represent the derived allele frequencies in Asian populations, against interpolated values of correlated dietary variables from Diet Model 1 for vitamin A status.

### Gene expression and allele-specific expression differences in genes associated with long-term dietary habits

Analysis of allele-specific expression (ASE) patterns for variants in the genes listed in Table S12 revealed that only two genes, *GHR* and *CAST*, contained variants that showed differences in expression. Allele-specific expression differences for all the individuals presented in Martin et al. (2014) as well as Pathan and Cambodia ethnic groups analyzed separately (see STAR Methods) showed lower expression in the Pathan than the Cambodian population (Figure S7). FPKM values showed differences in transcript levels for *GHR, SEMA3A* and *CAST* genes, with *CAST* showing more than two-fold FPKM values in Cambodia versus Pathan individuals (Figures S7 and S8). These results are indicative of the fact that with the availability of tissue-specific expression values in multiple populations, we can gain additional information on the functional attributes of the variants that track environmental variation.

## Discussion

Our findings underscore the importance of dietary transitions, and also provide a potential explanation for variation in disease risk which may differentially affect the viability (health and longevity) of individuals and families. For example, diet-related selection on genes related to olfaction and insulin production and sensitivity may help explain modern variation in taste preferences and diabetes etiology. Furthermore, knowing the historical availability of nutrients such as vitamin A, iron and zinc and its influence on genetic variation may help identify local ethnic groups/populations that are especially vulnerable to widespread modern-day deficiencies in the three micronutrients discussed. These results are especially compelling in light of the extensive genomic variation at a global and local scales that has been highlighted previously [21–23].

Of the highlighted genes and pathways in the present study, many are associated with type 2 diabetes and other metabolic disorders, or play a role in impaired metabolism or its dysregulation in relevant gene networks. Varying allele frequency patterns of the derived alleles under selection for each diet variable (Figures 4 and S9 and 10) may also suggest that each dietary factor had different impacts on fitness. For instance, the higher frequency of the derived allele of the *TCF7L2* variant (Figure 4) in East Asia, associated with protection from type 2 diabetes, may reflect an evolutionary trade-off. Here, vitamin A or a correlated dietary factor not represented in our model, created a selective pressure in East Asia that was advantageous in the past and protective against type 2 diabetes today. The next challenge will be to fine-map the variants that allow for efficient nutrient uptake, and understand the circumstances under which positive selection for each variant may have occurred.

It is important to note that any model, including ours, may be prone to systematic biases and errors. We failed to include several items of food that were significant components of the diets, were not endemic, or had only fragmentary physical evidence. We also did not take into account nutrient and genomic variation of ancient crops, or seasonal variability. Total caloric intake was also assumed to be constant through time, space and subsistence strategy. However, it is widely assumed that hunter-gatherers or those following a Paleolithic diet had higher caloric intake [24]. Furthermore, caloric density varies widely among the diets [25,26]. Finally, we did not include nutrient bioavailability, which is highly influenced by different food combinations, processing techniques, and evolving nutrient content [26,27].

Our study complements a growing body of literature showing that the more predictable and static external environmental factors such as elevation, climate and disease affect human evolution, by showing that dietary factors also drove genetic adaptations [28–31]. We have provided data that suggests regional adaptations to Neolithic diets which varied in nutrient profiles. Humans have shaped their own food environment over millennia, through a broadening range of food processing techniques and through domestication and the construction of agricultural systems [32–34]. These changes have left signatures on the genomes of different populations, and may have powerful implications for disparities in disease etiology as well as aging and life expectancy among historical and contemporary human populations [35]. In order to better understand these processes, it is necessary to collect detailed data on spatio-temporal variation in diet and the shift in agricultural production over millennia. These efforts would provide insights not only into the past human evolutionary genetic processes, but also regarding current problems of health and diseases in contemporary human populations.

## STAR Methods

## KEY RESOURCES TABLE

| REAGENT or RESOURCE | SOURCE | IDENTIFIER |
| --- | --- | --- |
| Deposited Data |  |  |
| Model 1 diet crops and timeline | Larson et al. (2014) | DOI:<br>10.1073/pnas.1323964111 |
| USDA food Composition databases | https://ndb.nal.usda.gov/ndb/ | N/A |
| Glycemic Index database | http://www.glycemicindex.com | N/A |
| Diogenes Glycemic Index database | http://www.diogenes-eu.org/GI-Database/Default.htm | N/A |
| Human Genome Diversity Panel (HGDP) | Metspalu et al. (2011) | DOI:<br>10.1016/j.ajhg.2011.11.010 |
| Genotype data – South Asia | G. Chaubey et al. (2011) | DOI:<br>10.1093/molbev/msq288 |
| Genotype data – South Asia | Behar et al. (2010) | DOI: 10.1038/nature09103 |
| HapMap 3 | HapMap Consortium | doi: 10.1038/nature09298 |
| HGDP genotype expression results | Martin et al. 2014 | DOI:<br>10.1371/journal.pgen.1004549 |
| Software and Algorithms |  |  |
| ADMIXTURE | Alexander et al. (2009) | DOI: [10.1101/gr.094052.109](https://doi.org/10.1101/gr.094052.109) |
| BayeScEnv | Villemereuil (2015) | DOI: 10.1111/2041-210X.12418 |
| BayeScan | Fischer et al. (2011), Foll and Gaggiotti (2008) | DOI: 10.1111/j.1365-294X.2011.05015.x, DOI: 10.1534/genetics.108.092221 |
| Baypass | Gautier (2015) | DOI: 10.1534/genetics.115.181453 |
| BEDtools | [Quinlan and Hall (2010)](https://doi.org/10.1093/bioinformatics/btq033) | DOI: 10.1093/bioinformatics/btq033 |
| Ensembl Variant Effect Predictor | http://www.ensembl.org/info/docs/tools/vep/index.html | DOI: 10.1186/s13059-016-0974-4 |
| GREAT | great.stanford.edu | DOI: 10.1038/nbt.1630 |
| LiftOver | https://genome.ucsc.edu/ | N/A |
| PLINK | http://zzz.bwh.harvard.edu/plink/ | DOI: 10.1086/519795 |
| traseR | Chen and Lin (2016) | DOI: 10.1093/bioinformatics/btv741 |
| Varanto | http://bioinformatics.uef.fi/varanto/ | DOI: [10.1093/bioinformatics/btz046](https://doi.org/10.1093/bioinformatics/btz046) |

### CONTACT FOR REAGENT AND RESOURCE SHARING

Further information and request for resources should be directed to the Lead Contact, Srilakshmi Raj.

## METHODS DETAILS

### Construction of dietary models from archaeological data

Archaeological data were used in two ways to estimate past dietary habits. First, we used the data displayed in Figure 2 in Larson et al. (2014), to account for the major domesticated species in Southwest Asia, South Asia and East Asia. Although this approach has excluded minor domesticates, it provides an empirically based simplification of regional variation in diet through time. We used 9000 years before present as a start point to infer dietary changes from agriculture and then added additional species diversity to regional diets over time. We assigned equal weights to all the dietary components and assumed they added up to 100% of the diet since the time of introduction (Table S1-S8). We took the start of the red bar in that figure (Larson et al. (2014) as an indicator of the time from which that organism was domesticated and added to the diet in that region.

To convert the archaeological data into a diet metric, we took information on the composition of micronutrients vitamin A, zinc, iron, omega-3, omega-6 and macronutrients protein, carbohydrates, and lipids for the cooked versions of the items in Figure 2 of Larson et al. (2014), coupled to the time parameter, and summed the resulting numbers to estimate the amount of each macro and micronutrient on the diet[36].

We constructed a second model to incorporate information on current population lifestyles (e.g. pastoralists, foragers and agriculturalists). We included hunter-gatherer diets as a proportion of the diet contributed by each item based on inference from archaeological evidence, and included hunter-gatherer diets as a proportion of the diet during the 9000 year time span. The second method of incorporating archaeological data to infer past diets differed from the first by: 1) varying the percentages of each item in the diet through time; 2) including a portion of hunter-gatherer diet; 3) differentiating diets of agricultural, pastoralist and cultivator populations; 4) varying food processing techniques; 5) using Glycemic Index (GI) and carbohydrate content as metrics for diet; and 6) using food substitutes for portions of the diet that were consumed and had similar glycemic index and carbohydrate content. GI and grams of carbohydrate per serving size were obtained from http://diogenes.s24.net/ and http://www.glycemicindex.com/[37–39]. These archaeology-based models were constructed based on both a literature search, the expertise of co-author Dorian Fuller, and knowledge of population habitats from Dr. G. Chaubey (pers. comm) (Table S9-S11).

Please see Supplementary Methods for additional details and references.

### Genetic Data

Genotype data from 506,306 SNPs from 1898 individuals belonging to 94 distinct global populations drawn from six published sources were filtered down to 29 populations from Asia for diet-based analyses (Tables S2,S3)[40–45]. We used PLINK to filter the data, using default settings for missingness, including a minor allele frequency threshold of 0.01[46]. Some of the South Asian populations represent composites of castes and tribes. The geographic coordinates were averaged across the samples included in each population (Table S11).

We used the locations of individuals from 29 populations with genome-wide SNP genotype data available to provide reference points for our dietary model (Table S10, Figure S1). The populations were assumed to have resided within a 500 km radius of the site over the millenia, for Baypass analysis. The method ADMIXTURE and information on ethnic background were used to group the 29 populations into 4 ethnic categories (Figure S1). These populations included six different subsistence strategies, belonged to six different countries and 4 ancestral groups (Figure S1, Table S10). For BayeScan and BayeScEnv analysis, populations were assumed to have resided within the entire regions captured by the ADMIXTURE groupings, over the millenia.

For the gene expression-based analyses in Figures S7 and S8, we included published data on whole transcriptome expression information in blood from the same Cambodia and Pathan individuals in this manuscript[47].

## QUANTIFICATION AND STATISTICAL ANALYSIS

### Detecting natural selection based on the dietary models

Three Bayesian methods, BayeScan, BayeScEnv and Baypass were used to detect differences in allele frequencies between populations, and to assess the significance of their correlations with environmental factors[48–51]. For the method BayPass, we specified an auxiliary model that assumed non-independence between environmental factors. All 9 dietary variables from both models were included in the same analysis. Further, we used the ADMIXTURE method to specify the population groups for the BayeScan analysis, and ascribing populations to archaeological data for the BayeScEnv analysis.

Since BayeScan relies on user-specified population groups, we used the software ADMIXTURE to determine genetic relatedness among the 94 populations studied[52]. We randomly sampled 20% of the genome-wide set of markers in the Asian subset of populations using PLINK, and tested a range of K values to determine the optimal number of population groups. We also used an option within ADMIXTURE to determine the cross-validation error attributed to each value of K. Subsequently, we plotted the estimated ancestry fractions using DISTRUCT software (Figure S1)[53]. A combination of the ancestral groupings determined by ADMIXTURE, and geographic location of populations were used to make population groupings for BayeScan analysis.

To assess significance for each dietary variable we took the top 1% of BayPass results and BayeScEnv results with q-values < 0.01. We also took BayeScan results with q-values < 1 × 10^−5^ and the highest Bayes Factor values. We used the Liftover program in the UCSC genome browser to convert from NCBI36 (hg18) to NCBI37 (hg19) genomic positions[54]. To obtain functional annotations of SNPs, we entered the updated rs numbers into the Ensembl Variant Effect Predictor online tool (http://www.ensembl.org/Homo_sapiens/Tools/VEP)[55] to obtain genome positions and annotations for these SNPs, including functional class and PolyPhen and SIFT score. All figures were made using the ggplot2 package in R (v. 3.5.1). Significance of enrichment within each functional category, PolyPhen and SIFT scores were assessed using Varanto (http://bioinformatics.uef.fi/varanto/), against the reference Ilumina Human660W-Quad [56].

The Gene Ontology program GREAT was used to identify categories under “Biological Process” that were significantly enriched in these signals[57]. To provide input regions for GREAT we ascribed to each SNP a short segment spanning ±500 bp. Then we used BEDtools to merge consecutive regions spaced less than 1000 bp apart into larger regions for GREAT input[57,58].

The program traseR[19] was used to perform enrichment analyses of selection results in genome-wide association studies, with the latter data extracted from the NIH NHGRI-EBI GWA database (https://www.ebi.ac.uk/gwas/). Default settings were used, including a binomial test for enrichment.

### Analysis of expression data

Correlations among long-term dietary habits and genetic variation that influence phenotypic variation suggests that the expression and function of these genes may vary contextually among different parts of Asia. For this analysis, we used published data on whole transcriptome expression information in blood taken from the same Human Genome Diversity Panel individuals included in this manuscript[47]. Of the populations included in Martin et al. (2014) and the present study, only two, Cambodia and Pathan, overlapped. Each of these were considered representative of the East Asia and West Asia populations. These data provided the allele-specific expression (ASE) and Fragments Per Kilobase of transcript per Million mapped reads (FPKM) results in Figures S7 and S8.

