## Supplementary Figures, Methods and Tables for "Reconstruction of nine thousand years of agriculture-based diet and impact on human genetic diversity in Asia"

### Supplementary Methods

#### Reconstruction of past dietary habits based on archeological data - diet model 1

We estimated dietary habits across Asia over a 9000 year time span using the archaeological data presented in Figure 2 of Larson et al. (2014)[1]. We used their indicators of the latest time by which domestication occurred, and used a time span of 9000 years to represent a timeframe within which agricultural food production was practiced. For those plants and animals that were domesticated prior to 9000 years ago, those earlier dates were included in our model as well (Table S1-S8).

We used the ‘Southwest Asia’, ‘South Asia’ and ‘East Asia’ categories in Figure 2 of Larson et al. (2014) to make inferences of diet over this time span. Regions were necessarily broad to allow comparison of genetic data, although we recognize this may merge smaller regions that archaeologists might regard as distinct. We have taken Pakistan populations as representative of Southwest Asia since archaeologically the agriculture in this region was focused on species deriving from further west such as sheep, goat, cattle, wheat and barley[2], whereas South Asia represents populations from regions that experienced earlier consumption of rice, native millets and pulses. Because the items in this table represented domestication of some of the most important plants and animals in their respective geographic regions, we assumed that these items were consumed in equal amounts, and together added up to 100% of the diets of all individuals and populations in these regions, starting from the latest time of domestication. All items in this figure were assumed to be consumed as is, with the exception of ‘Bactrian camel’, ‘silk worm’ and ‘tree cotton’ as objects of dietary consumption. We used macro and micronutrient composition of modern-day equivalents of these species to extrapolate the dietary habits of the populations in these regions[3]. The macronutrients included: carbohydrates, proteins and lipids, the latter of which we also divided into omega-3 and omega-6 fatty acids, due to their essential role in the human diet as well as in health and disease[4–6]. Because vitamin A, zinc and iron deficiencies are common nutrient deficiencies, especially in South Asia, contributing to stunting[7–9], we also included these values in our model. All values were provided as g/100g of dry weight. We multiplied these values by the time of exposure relative to 9000 years, and added these values for all compounds to obtain a dietary metric for each of the three regional groups (Figure 2, Figure 3, Table S1-S8).

As carbohydrate consumption was a prominent feature in the Agricultural Transition, we used glycemic index (a measurement of the increase in blood glucose after food consumption), and carbohydrates contributed by the food item to the total diet. GI and carbohydrate content provide useful indices of health effects of foods, because they may be related to T2D risk and related health complications. Low GI foods have been shown to provide some protection to T2D, or high GI food and high carbohydrate content in the diet with increased risk to T2D[10–14], although this has been contested[15]. Note that T2D is also a modern epidemic, unlikely to have influenced health over evolutionary time scales in humans[16].

We assigned the South Asia, Southwest Asia and East Asia dietary models to the populations belonging to the ADMIXTURE categories 1, 2 and 3, respectively. The ADMIXTURE category 4 populations were assigned to Southwest Asia.

**Reconstruction of past dietary habits based on archeological data - diet model 2**
**Assumptions:**
For the second diet model, we used a seven-step process to associate archaeological evidence of crop and animal domestication with current populations (Figure S1, Table S9). We made the following assumptions: a) all populations studied resided within a 500 km radius from their current location over the last 9,000 years ( ~360 generations); b) caloric intake has remained approximately similar across populations and through time; c) the dietary models represented the complete diet for the populations during the time slice, and d) assuming Wright's island model, there were few enough migration, admixture, mutation events through time in a given area that they could be ignored. Subsequently, the 29 Asian populations with genotype data were associated with archaeological evidence of crop domestication within a 500 km radius of an archaeological site (Figure S1). Furthermore, we estimated an almost uniform hunter-gatherer diet for a baseline period to the advent of agriculture, with only modest regional differences. However, it is likely that hunter-gatherer societies varied among different eco-regions as is true in ethnographic hunter-gatherers[17]. In addition, we did not include certain food items that were strictly seasonal, such as honey, although it is suggested to have been important in hominin diet[18,19].

**Dietary components:**
We used two dietary components to quantify variation in diet: carbohydrate content and glycemic index. Both were obtained from the databases http://www.glycemicindex.com/ and http://www.diogenes-eu.org/GI-Database/Default.htm . Food items taken from the Diogenes database have an additional item label listed next to the food item in Supplementary Table 9. For example, "carp" is listed in the table without any other label, but beside "raw" is the additional label "UK-2533", form the Diogenes database. Carbohydrate content was expressed as grams of carbohydrate per 100g of prepared food. The "carbs contributed by food item" reflects the carbohydrate content in the entire dietary period if the whole estimated diet for that population and time period ("time slice") were a single 100g serving. We also calculated glycemic index for each time slice as if it were a single meal, accounting for carbohydrate content (Dr. Susan Jebb, pers. comm.).

The diet models were constructed using items native to the geographic region, or where unavailable, closest proxies available in the Glycemic Index databases[20,21]. The primary departure from use of native crops across regions was the translocation of crops across regions, such as the introduction of wheat into China or India or African sorghum into India[22,23]. However, despite early introductions some of these crops may not have become dietarily important until much more recently, such as around 2500 YBP for wheat in central China or sorghum to the diet of the South India caste groups, as it became an important part of their diet since 1500 YBP[24]. To define the earliest time for widespread consumption of each item, we subtracted 500 to 1000 years off the earliest appearance of the item in the archaeological record. For example, the earliest evidence for rice farming in East Asia appears 8000 YBP, but 3000 YBP in Japan[25]. Therefore, Han and She populations were estimated to widely consume rice from 7000 YBP and Japan populations from 3000 YBP (Table S9). We divided 9000 years into 1000 year segments of time to incorporate the different times that each item became included in the diet. The earlier time slices had larger 'gaps' unoccupied by domesticated food items introduced later in time. We used the average estimation for the preceding time slice to fill these gaps.

**Supplementary Table 1.** Model of lipid consumption over 9000 years in Asia, to construct Diet Model 1. The Total columns for each geographic region represents the proportion of lipids/100g contributed by the specific crop in each geographic region, respectively.

| | LIPIDS | | | | | | | | |
|---|---|---|---|---|---|---|---|---|---|
| | Southwest Asia | | | South Asia | | | East Asia | | |
| | Lipids (g/100g) | Proportion of years (out of 9000) of domestication | Total | Lipids (g/100g) | Proportion of years (out of 9000) of domestication | Total | Lipids (g/100g) | Proportion of years (out of 9000) of domestication | Total |
| Wheat | 1.30 | 1.00 | 1.30 | 0.00 | 0.00 | 0.00 | 0.00 | 0.00 | 0.00 |
| Barley | 1.20 | 1.00 | 1.20 | 0.00 | 0.00 | 0.00 | 0.00 | 0.00 | 0.00 |
| Lentil | 1.10 | 1.00 | 1.10 | 0.00 | 0.00 | 0.00 | 0.00 | 0.00 | 0.00 |
| Pea | 0.40 | 0.94 | 0.38 | 0.00 | 0.00 | 0.00 | 0.00 | 0.00 | 0.00 |
| Chickpea | 6.00 | 0.92 | 5.50 | 0.00 | 0.00 | 0.00 | 0.00 | 0.00 | 0.00 |
| Broadbean | 1.50 | 1.17 | 1.75 | 0.00 | 0.00 | 0.00 | 0.00 | 0.00 | 0.00 |
| Flax | 42.20 | 1.06 | 44.54 | 0.00 | 0.00 | 0.00 | 0.00 | 0.00 | 0.00 |
| Olive | 15.30 | 0.67 | 10.20 | 0.00 | 0.00 | 0.00 | 0.00 | 0.00 | 0.00 |
| Sheep | 15.00 | 0.89 | 13.33 | 0.00 | 0.00 | 0.00 | 0.00 | 0.00 | 0.00 |
| Goat | 2.30 | 0.89 | 2.04 | 0.00 | 0.00 | 0.00 | 0.00 | 0.00 | 0.00 |
| Cattle (taurine) | 22.50 | 0.89 | 20.00 | 0.00 | 0.00 | 0.00 | 0.00 | 0.00 | 0.00 |
| Cat | 0.00 | 0.44 | 0.00 | 0.00 | 0.00 | 0.00 | 0.00 | 0.00 | 0.00 |
| Tree Cotton | | 0.00 | 0.00 | | 0.50 | 0.00 | | 0.00 | 0.00 |
| Rice (indica) | 0.00 | 0.00 | 0.00 | 1.10 | 0.28 | 0.31 | 0.00 | 0.00 | 0.00 |
| Little Millet | 0.00 | 0.00 | 0.00 | 4.20 | 0.50 | 2.10 | 0.00 | 0.00 | 0.00 |
| Browntop Millet | 0.00 | 0.00 | 0.00 | 4.20 | 0.44 | 1.87 | 0.00 | 0.00 | 0.00 |
| Mungbean | 0.00 | 0.00 | 0.00 | 1.20 | 0.33 | 0.40 | 0.00 | 0.00 | 0.00 |
| Pigeonpea | 0.00 | 0.00 | 0.00 | 1.60 | 0.39 | 0.62 | 0.00 | 0.00 | 0.00 |
| Cattle (zebu) | 0.00 | 0.00 | 0.00 | 22.50 | 0.72 | 16.25 | 0.00 | 0.00 | 0.00 |
| Water buffalo | 0.00 | 0.00 | 0.00 | 1.40 | 0.50 | 0.70 | 0.00 | 0.00 | 0.00 |
| Broomcorn millet | 0.00 | 0.00 | 0.00 | 0.00 | 0.00 | 0.00 | 4.20 | 0.89 | 3.73 |
| Foxtail millet | 0.00 | 0.00 | 0.00 | 0.00 | 0.00 | 0.00 | 4.20 | 0.83 | 3.50 |
| Rice (japonica) | 0.00 | 0.00 | 0.00 | 0.00 | 0.00 | 0.00 | 1.10 | 0.56 | 0.61 |
| Soybean | 0.00 | 0.00 | 0.00 | 0.00 | 0.00 | 0.00 | 19.90 | 0.39 | 7.74 |
| Ramie | 0.00 | 0.00 | 0.00 | 0.00 | 0.00 | 0.00 | 0.30 | 0.58 | 0.17 |
| Melon | 0.00 | 0.00 | 0.00 | 0.00 | 0.00 | 0.00 | 0.10 | 0.39 | 0.04 |
| Pig | 71.00 | 1.00 | 71.00 | 0.00 | 0.00 | 0.00 | 71.00 | 0.67 | 47.33 |
| Silkworm | 0.00 | 0.00 | 0.00 | 0.00 | 0.00 | 0.00 | 8.76 | 0.64 | 5.60 |
| Yak | 0.00 | 0.00 | 0.00 | 0.00 | 0.00 | 0.00 | 7.08 | 0.47 | 3.34 |
| Horse | 0.00 | 0.00 | 0.00 | 0.00 | 0.00 | 0.00 | 4.60 | 0.44 | 2.04 |

| | | | | | | | | | |
|---|---|---|---|---|---|---|---|---|---|
| Bactrian Camel | 0.00 | 0.00 | 0.00 | 0.00 | 0.00 | 0.00 | 1.90 | 0.50 | 0.95 |
| Duck | 0.00 | 0.00 | 0.00 | 0.00 | 0.00 | 0.00 | 39.30 | 0.11 | 4.37 |
| Chicken | 0.00 | 0.00 | 0.00 | 0.00 | 0.00 | 0.00 | 8.10 | 0.44 | 3.60 |
| TOTAL (Average) | | | 5.22 | | | 0.67 | | | 2.52 |

**Supplementary Table 2.** Model of iron consumption over 9000 years in Asia, to construct Diet Model 1. The Total columns for each geographic region represents the proportion of iron/100g contributed by the specific crop in each geographic region, respectively.

| IRON | | | | | | | | | |
|---|---|---|---|---|---|---|---|---|---|
| | Southwest Asia | | | South Asia | | | East Asia | | |
| | iron (g/100g) | Proportion of years (out of 9000) of domestication | Total | Iron (g/100g) | Proportion of years (out of 9000) of domestication | Total | iron (g/100g) | Proportion of years (out of 9000) of domestication | Total |
| Wheat | 0.0021 | 1.0000 | 0.0021 | 0.0000 | 0.0000 | 0.0000 | 0.0000 | 0.0000 | 0.0000 |
| Barley | 0.0025 | 1.0000 | 0.0025 | 0.0000 | 0.0000 | 0.0000 | 0.0000 | 0.0000 | 0.0000 |
| Lentil | 0.0075 | 1.0000 | 0.0075 | 0.0000 | 0.0000 | 0.0000 | 0.0000 | 0.0000 | 0.0000 |
| Pea | 0.0015 | 0.9444 | 0.0014 | 0.0000 | 0.0000 | 0.0000 | 0.0000 | 0.0000 | 0.0000 |
| Chickpea | 0.0062 | 0.9167 | 0.0057 | 0.0000 | 0.0000 | 0.0000 | 0.0000 | 0.0000 | 0.0000 |
| Broadbean | 0.0067 | 1.1667 | 0.0078 | 0.0000 | 0.0000 | 0.0000 | 0.0000 | 0.0000 | 0.0000 |
| Flax | 0.0057 | 1.0556 | 0.0060 | 0.0000 | 0.0000 | 0.0000 | 0.0000 | 0.0000 | 0.0000 |
| Olive | 0.0005 | 0.6667 | 0.0003 | 0.0000 | 0.0000 | 0.0000 | 0.0000 | 0.0000 | 0.0000 |
| Sheep | 0.0011 | 0.8889 | 0.0010 | 0.0000 | 0.0000 | 0.0000 | 0.0000 | 0.0000 | 0.0000 |
| Goat | 0.0028 | 0.8889 | 0.0025 | 0.0000 | 0.0000 | 0.0000 | 0.0000 | 0.0000 | 0.0000 |
| Cattle (taurine) | 0.0019 | 0.8889 | 0.0017 | 0.0000 | 0.0000 | 0.0000 | 0.0000 | 0.0000 | 0.0000 |
| Cat | 0.0000 | 0.4444 | 0.0000 | 0.0000 | 0.0000 | 0.0000 | 0.0000 | 0.0000 | 0.0000 |
| Tree Cotton | 0.0000 | 0.0000 | 0.0000 | 0.0000 | 0.5000 | 0.0000 | 0.0000 | 0.0000 | 0.0000 |

| | | | | | | | | | |
|---|---|---|---|---|---|---|---|---|---|
| Rice (indica) | 0.0000 | 0.0000 | 0.0000 | 0.0020 | 0.2778 | 0.0006 | 0.0000 | 0.0000 | 0.0000 |
| Little Millet | 0.0000 | 0.0000 | 0.0000 | 0.0030 | 0.5000 | 0.0015 | 0.0000 | 0.0000 | 0.0000 |
| Browntop Millet | 0.0000 | 0.0000 | 0.0000 | 0.0030 | 0.4444 | 0.0013 | 0.0000 | 0.0000 | 0.0000 |
| Mungbean | 0.0000 | 0.0000 | 0.0000 | 0.0067 | 0.3333 | 0.0022 | 0.0000 | 0.0000 | 0.0000 |
| Pigeonpea | 0.0000 | 0.0000 | 0.0000 | 0.0016 | 0.3889 | 0.0006 | 0.0000 | 0.0000 | 0.0000 |
| Cattle (zebu) | 0.0000 | 0.0000 | 0.0000 | 0.0019 | 0.7222 | 0.0014 | 0.0000 | 0.0000 | 0.0000 |
| Water buffalo | 0.0000 | 0.0000 | 0.0000 | 0.0016 | 0.5000 | 0.0008 | 0.0000 | 0.0000 | 0.0000 |
| Broomcorn millet | 0.0000 | 0.0000 | 0.0000 | 0.0000 | 0.0000 | 0.0000 | 0.0030 | 0.8889 | 0.0027 |
| Foxtail millet | 0.0000 | 0.0000 | 0.0000 | 0.0000 | 0.0000 | 0.0000 | 0.0030 | 0.8333 | 0.0025 |
| Rice (japonica) | 0.0000 | 0.0000 | 0.0000 | 0.0000 | 0.0000 | 0.0000 | 0.0020 | 0.5556 | 0.0011 |
| Soybean | 0.0000 | 0.0000 | 0.0000 | 0.0000 | 0.0000 | 0.0000 | 0.0157 | 0.3889 | 0.0061 |
| Ramie | 0.0000 | 0.0000 | 0.0000 | 0.0000 | 0.0000 | 0.0000 | 0.0048 | 0.5833 | 0.0028 |
| Melon | 0.0000 | 0.0000 | 0.0000 | 0.0000 | 0.0000 | 0.0000 | 0.0002 | 0.3889 | 0.0001 |
| Pig | 0.0004 | 1.0000 | 0.0004 | 0.0000 | 0.0000 | 0.0000 | 0.0004 | 0.6667 | 0.0003 |
| Silkworm | 0.0000 | 0.0000 | 0.0000 | 0.0000 | 0.0000 | 0.0000 | 0.0095 | 0.6389 | 0.0061 |
| Yak | 0.0000 | 0.0000 | 0.0000 | 0.0000 | 0.0000 | 0.0000 | 0.0027 | 0.4722 | 0.0013 |
| Horse | 0.0000 | 0.0000 | 0.0000 | 0.0000 | 0.0000 | 0.0000 | 0.0038 | 0.4444 | 0.0017 |
| Bactrian Camel | 0.0000 | 0.0000 | 0.0000 | 0.0000 | 0.0000 | 0.0000 | 0.0001 | 0.5000 | 0.0001 |
| Duck | 0.0000 | 0.0000 | 0.0000 | 0.0000 | 0.0000 | 0.0000 | 0.0024 | 0.1111 | 0.0003 |
| Chicken | 0.0000 | 0.0000 | 0.0000 | 0.0000 | 0.0000 | 0.0000 | 0.0008 | 0.4444 | 0.0004 |
| TOTAL (Average) | | | 0.0012 | | | 0.0003 | | | 0.0008 |

**Supplementary Table 3.** Model of zinc consumption over 9000 years in Asia, to construct Diet Model 1. The Total columns for each geographic region represents the proportion of zinc/100g contributed by the specific crop in each geographic region, respectively.

| ZINC | | | |
|---|---|---|---|
| | Southwest Asia | South Asia | East Asia |

| | zinc (g/100g) | Proportion of years (out of 9000) of domestication | Total | zinc (g/100g) | Proportion of years (out of 9000) of domestication | Total | zinc (g/100g) | Proportion of years (out of 9000) of domestication | Total |
|---|---|---|---|---|---|---|---|---|---|
| Wheat | 0.0017 | 1.0000 | 0.0017 | 0.0000 | 0.0000 | 0.0000 | 0.0000 | 0.0000 | 0.0000 |
| Barley | 0.0021 | 1.0000 | 0.0021 | 0.0000 | 0.0000 | 0.0000 | 0.0000 | 0.0000 | 0.0000 |
| Lentil | 0.0048 | 1.0000 | 0.0048 | 0.0000 | 0.0000 | 0.0000 | 0.0000 | 0.0000 | 0.0000 |
| Pea | 0.0012 | 0.9444 | 0.0011 | 0.0000 | 0.0000 | 0.0000 | 0.0000 | 0.0000 | 0.0000 |
| Chickpea | 0.0034 | 0.9167 | 0.0031 | 0.0000 | 0.0000 | 0.0000 | 0.0000 | 0.0000 | 0.0000 |
| Broadbean | 0.0031 | 1.1667 | 0.0036 | 0.0000 | 0.0000 | 0.0000 | 0.0000 | 0.0000 | 0.0000 |
| Flax | 0.0043 | 1.0556 | 0.0045 | 0.0000 | 0.0000 | 0.0000 | 0.0000 | 0.0000 | 0.0000 |
| Olive | 0.0000 | 0.6667 | 0.0000 | 0.0000 | 0.0000 | 0.0000 | 0.0000 | 0.0000 | 0.0000 |
| Sheep | 0.0030 | 0.8889 | 0.0026 | 0.0000 | 0.0000 | 0.0000 | 0.0000 | 0.0000 | 0.0000 |
| Goat | 0.0040 | 0.8889 | 0.0036 | 0.0000 | 0.0000 | 0.0000 | 0.0000 | 0.0000 | 0.0000 |
| Cattle (taurine) | 0.0036 | 0.8889 | 0.0032 | 0.0000 | 0.0000 | 0.0000 | 0.0000 | 0.0000 | 0.0000 |
| Cat | 0.0000 | 0.4444 | 0.0000 | 0.0000 | 0.0000 | 0.0000 | 0.0000 | 0.0000 | 0.0000 |
| Tree Cotton | 0.0000 | 0.0000 | 0.0000 | 0.0000 | 0.5000 | 0.0000 | 0.0000 | 0.0000 | 0.0000 |
| Rice (indica) | 0.0000 | 0.0000 | 0.0000 | 0.0060 | 0.2778 | 0.0017 | 0.0000 | 0.0000 | 0.0000 |
| Little Millet | 0.0000 | 0.0000 | 0.0000 | 0.0017 | 0.5000 | 0.0009 | 0.0000 | 0.0000 | 0.0000 |
| Browntop Millet | 0.0000 | 0.0000 | 0.0000 | 0.0017 | 0.4444 | 0.0008 | 0.0000 | 0.0000 | 0.0000 |
| Mungbean | 0.0000 | 0.0000 | 0.0000 | 0.0027 | 0.3333 | 0.0009 | 0.0000 | 0.0000 | 0.0000 |
| Pigeonpea | 0.0000 | 0.0000 | 0.0000 | 0.0010 | 0.3889 | 0.0004 | 0.0000 | 0.0000 | 0.0000 |
| Cattle (zebu) | 0.0000 | 0.0000 | 0.0000 | 0.0036 | 0.7222 | 0.0026 | 0.0000 | 0.0000 | 0.0000 |
| Water buffalo | 0.0000 | 0.0000 | 0.0000 | 0.0019 | 0.5000 | 0.0010 | 0.0000 | 0.0000 | 0.0000 |
| Broomcorn millet | 0.0000 | 0.0000 | 0.0000 | 0.0000 | 0.0000 | 0.0000 | 0.0017 | 0.8889 | 0.0015 |
| Foxtail millet | 0.0000 | 0.0000 | 0.0000 | 0.0000 | 0.0000 | 0.0000 | 0.0017 | 0.8333 | 0.0014 |
| Rice (japonica) | 0.0000 | 0.0000 | 0.0000 | 0.0000 | 0.0000 | 0.0000 | 0.0060 | 0.5556 | 0.0033 |
| Soybean | 0.0000 | 0.0000 | 0.0000 | 0.0000 | 0.0000 | 0.0000 | 0.0049 | 0.3889 | 0.0019 |
| Ramie | 0.0000 | 0.0000 | 0.0000 | 0.0000 | 0.0000 | 0.0000 | 0.0008 | 0.5833 | 0.0005 |
| Melon | 0.0000 | 0.0000 | 0.0000 | 0.0000 | 0.0000 | 0.0000 | 0.0001 | 0.3889 | 0.0000 |
| Pig | 0.0007 | 1.0000 | 0.0007 | 0.0000 | 0.0000 | 0.0000 | 0.0007 | 0.6667 | 0.0005 |
| Silkworm | 0.0000 | 0.0000 | 0.0000 | 0.0000 | 0.0000 | 0.0000 | 0.0178 | 0.6389 | 0.0113 |
| Yak | 0.0000 | 0.0000 | 0.0000 | 0.0000 | 0.0000 | 0.0000 | 0.0000 | 0.4722 | 0.0000 |
| Horse | 0.0000 | 0.0000 | 0.0000 | 0.0000 | 0.0000 | 0.0000 | 0.0029 | 0.4444 | 0.0013 |
| Bactrian Camel | 0.0000 | 0.0000 | 0.0000 | 0.0000 | 0.0000 | 0.0000 | 0.0003 | 0.5000 | 0.0002 |
| Duck | 0.0000 | 0.0000 | 0.0000 | 0.0000 | 0.0000 | 0.0000 | 0.0014 | 0.1111 | 0.0002 |
| Chicken | 0.0000 | 0.0000 | 0.0000 | 0.0000 | 0.0000 | 0.0000 | 0.0015 | 0.4444 | 0.0007 |
| TOTAL (Average) | | | 0.0009 | | | 0.0002 | | | 0.0007 |

**Supplementary Table 4.** Model of vitamin A consumption over 9000 years in Asia, to construct Diet Model 1. The Total columns for each geographic region represents the proportion of vitamin A (IU) contributed by the specific crop in each geographic region, respectively.

| VITAMIN A | | | | | | | | | |
|---|---|---|---|---|---|---|---|---|---|
| | Southwest Asia | | | South Asia | | | East Asia | | |
| | IU | Proportion of years (out of 9000) of domestication | Total | IU | Proportion of years (out of 9000) of domestication | Total | IU | Proportion of years (out of 9000) of domestication | Total |
| Wheat | 0.00 | 1.00 | 0.00 | 0.00 | 0.00 | 0.00 | 0.00 | 0.00 | 0.00 |
| Barley | 22.00 | 1.00 | 22.00 | 0.00 | 0.00 | 0.00 | 0.00 | 0.00 | 0.00 |
| Lentil | 39.00 | 1.00 | 39.00 | 0.00 | 0.00 | 0.00 | 0.00 | 0.00 | 0.00 |
| Pea | 765.00 | 0.94 | 722.50 | 0.00 | 0.00 | 0.00 | 0.00 | 0.00 | 0.00 |
| Chickpea | 67.00 | 0.92 | 61.42 | 0.00 | 0.00 | 0.00 | 0.00 | 0.00 | 0.00 |
| Broadbean | 53.00 | 1.17 | 61.83 | 0.00 | 0.00 | 0.00 | 0.00 | 0.00 | 0.00 |
| Flax | 0.00 | 1.06 | 0.00 | 0.00 | 0.00 | 0.00 | 0.00 | 0.00 | 0.00 |
| Olive | 393.00 | 0.67 | 262.00 | 0.00 | 0.00 | 0.00 | 0.00 | 0.00 | 0.00 |
| Sheep | 0.00 | 0.89 | 0.00 | 0.00 | 0.00 | 0.00 | 0.00 | 0.00 | 0.00 |
| Goat | 0.00 | 0.89 | 0.00 | 0.00 | 0.00 | 0.00 | 0.00 | 0.00 | 0.00 |
| Cattle(taurine) | 0.00 | 0.89 | 0.00 | 0.00 | 0.00 | 0.00 | 0.00 | 0.00 | 0.00 |
| Cat | 0.00 | 0.44 | 0.00 | 0.00 | 0.00 | 0.00 | 0.00 | 0.00 | 0.00 |
| Tree Cotton | 0.00 | 0.00 | 0.00 | 0.00 | 0.50 | 0.00 | 0.00 | 0.00 | 0.00 |
| Rice (indica) | 0.00 | 0.00 | 0.00 | 19.00 | 0.28 | 5.28 | 0.00 | 0.00 | 0.00 |
| Little Millet | 0.00 | 0.00 | 0.00 | 0.00 | 0.50 | 0.00 | 0.00 | 0.00 | 0.00 |
| Browntop Millet | 0.00 | 0.00 | 0.00 | 0.00 | 0.44 | 0.00 | 0.00 | 0.00 | 0.00 |
| Mungbean | 0.00 | 0.00 | 0.00 | 114.00 | 0.33 | 38.00 | 0.00 | 0.00 | 0.00 |
| Pigeonpea | 0.00 | 0.00 | 0.00 | 67.00 | 0.39 | 26.06 | 0.00 | 0.00 | 0.00 |
| Cattle (zebu) | 0.00 | 0.00 | 0.00 | 0.00 | 0.72 | 0.00 | 0.00 | 0.00 | 0.00 |
| Water buffalo | 0.00 | 0.00 | 0.00 | 0.00 | 0.50 | 0.00 | 0.00 | 0.00 | 0.00 |
| Broomcorn millet | 0.00 | 0.00 | 0.00 | 0.00 | 0.00 | 0.00 | 2.99 | 0.89 | 2.66 |
| Foxtail millet | 0.00 | 0.00 | 0.00 | 0.00 | 0.00 | 0.00 | 0.00 | 0.83 | 0.00 |
| Rice (japonica) | 0.00 | 0.00 | 0.00 | 0.00 | 0.00 | 0.00 | 19.00 | 0.56 | 10.56 |
| Soybean | 0.00 | 0.00 | 0.00 | 0.00 | 0.00 | 0.00 | 22.00 | 0.39 | 8.56 |
| Ramie | 0.00 | 0.00 | 0.00 | 0.00 | 0.00 | 0.00 | 5,559.00 | 0.58 | 3,242.75 |
| Melon | 0.00 | 0.00 | 0.00 | 0.00 | 0.00 | 0.00 | 50.00 | 0.39 | 19.44 |
| Pig | 81.00 | 1.00 | 81.00 | 0.00 | 0.00 | 0.00 | 81.00 | 0.67 | 54.00 |
| Silkworm | 0.00 | 0.00 | 0.00 | 0.00 | 0.00 | 0.00 | 0.00 | 0.64 | 0.00 |
| Yak | 0.00 | 0.00 | 0.00 | 0.00 | 0.00 | 0.00 | 0.00 | 0.47 | 0.00 |
| Horse | 0.00 | 0.00 | 0.00 | 0.00 | 0.00 | 0.00 | 0.00 | 0.44 | 0.00 |
| Bactrian Camel | 0.00 | 0.00 | 0.00 | | 0.00 | 0.00 | 83.40 | 0.50 | 41.70 |
| Duck | 0.00 | 0.00 | 0.00 | 0.00 | 0.00 | 0.00 | 168.00 | 0.11 | 18.66 |
| Chicken | 0.00 | 0.00 | 0.00 | 0.00 | 0.00 | 0.00 | 0.00 | 0.44 | 0.00 |
| TOTAL (Average) | | | 37.87 | | | 2.10 | | | 102.98 |

**Supplementary Table 5.** Model of omega-6 consumption over 9000 years in Asia, to construct Diet Model 1. The Total columns for each geographic region represents the proportion of omega-6 (g/100g) contributed by the specific crop in each geographic region, respectively.

| OMEGA 6 | | | | | | | | | |
|---|---|---|---|---|---|---|---|---|---|
| | Southwest Asia | | | South Asia | | | East Asia | | |
| | g/100g | Proportion of years (out of 9000) of domestication | Total | g/100g | Proportion of years (out of 9000) of domestication | Total | g/100g | Proportion of years (out of 9000) of domestication | Total |
| Wheat | 0.53 | 1.00 | 0.53 | 0.00 | 0.00 | 0.00 | 0.00 | 0.00 | 0.00 |
| Barley | 0.51 | 1.00 | 0.51 | 0.00 | 0.00 | 0.00 | 0.00 | 0.00 | 0.00 |
| Lentil | 0.40 | 1.00 | 0.40 | 0.00 | 0.00 | 0.00 | 0.00 | 0.00 | 0.00 |
| Pea | 0.15 | 0.94 | 0.14 | 0.00 | 0.00 | 0.00 | 0.00 | 0.00 | 0.00 |
| Chickpea | 2.59 | 0.92 | 2.38 | 0.00 | 0.00 | 0.00 | 0.00 | 0.00 | 0.00 |
| Broadbean | 0.58 | 1.17 | 0.68 | 0.00 | 0.00 | 0.00 | 0.00 | 0.00 | 0.00 |
| Flax | 5.91 | 1.06 | 6.24 | 0.00 | 0.00 | 0.00 | 0.00 | 0.00 | 0.00 |
| Olive | 1.22 | 0.67 | 0.81 | 0.00 | 0.00 | 0.00 | 0.00 | 0.00 | 0.00 |
| Sheep | 0.00 | 0.89 | 0.00 | 0.00 | 0.00 | 0.00 | 0.00 | 0.00 | 0.00 |
| Goat | 0.10 | 0.89 | 0.09 | 0.00 | 0.00 | 0.00 | 0.00 | 0.00 | 0.00 |
| Cattle(taurine) | 0.57 | 0.89 | 0.51 | 0.00 | 0.00 | 0.00 | 0.00 | 0.00 | 0.00 |
| Cat | 0.00 | 0.44 | 0.00 | 0.00 | 0.00 | 0.00 | 0.00 | 0.00 | 0.00 |
| Tree Cotton | 0.00 | 0.00 | 0.00 | 0.00 | 0.50 | 0.00 | 0.00 | 0.00 | 0.00 |
| Rice (indica) | 0.00 | 0.00 | 0.00 | 0.38 | 0.28 | 0.10 | 0.00 | 0.00 | 0.00 |
| Little Milet | 0.00 | 0.00 | 0.00 | 2.02 | 0.50 | 1.01 | 0.00 | 0.00 | 0.00 |
| Browntop Millet | 0.00 | 0.00 | 0.00 | 2.02 | 0.44 | 0.90 | 0.00 | 0.00 | 0.00 |
| Mungbean | 0.00 | 0.00 | 0.00 | 0.36 | 0.33 | 0.12 | 0.00 | 0.00 | 0.00 |
| Pigeonpea | 0.00 | 0.00 | 0.00 | 0.84 | 0.39 | 0.32 | 0.00 | 0.00 | 0.00 |
| Cattle (zebu) | 0.00 | 0.00 | 0.00 | 0.57 | 0.72 | 0.41 | 0.00 | 0.00 | 0.00 |
| Water buffalo | 0.00 | 0.00 | 0.00 | 0.16 | 0.50 | 0.08 | 0.00 | 0.00 | 0.00 |
| Broomcorn millet | 0.00 | 0.00 | 0.00 | 0.00 | 0.00 | 0.00 | 2.02 | 0.89 | 1.79 |
| Foxtail millet | 0.00 | 0.00 | 0.00 | 0.00 | 0.00 | 0.00 | 2.02 | 0.83 | 1.68 |
| Rice (japonica) | 0.00 | 0.00 | 0.00 | 0.00 | 0.00 | 0.00 | 0.38 | 0.56 | 0.21 |
| Soybean | 0.00 | 0.00 | 0.00 | 0.00 | 0.00 | 0.00 | 9.92 | 0.39 | 3.86 |
| Ramie | 0.00 | 0.00 | 0.00 | 0.00 | 0.00 | 0.00 | 0.12 | 0.58 | 0.07 |
| Melon | 0.00 | 0.00 | 0.00 | 0.00 | 0.00 | 0.00 | 0.03 | 0.39 | 0.01 |
| Pig | 9.31 | 1.00 | 9.31 | 0.00 | 0.00 | 0.00 | 9.31 | 0.67 | 6.21 |
| Silkworm | 0.00 | 0.00 | 0.00 | 0.00 | 0.00 | 0.00 | 0.00 | 0.64 | 0.00 |
| Yak | 0.00 | 0.00 | 0.00 | 0.00 | 0.00 | 0.00 | 0.00 | 0.47 | 0.00 |
| Horse | 0.00 | 0.00 | 0.00 | 0.00 | 0.00 | 0.00 | 0.29 | 0.44 | 0.13 |
| Bactrian Camel | 0.00 | 0.00 | 0.00 | 0.00 | 0.00 | 0.00 | 0.00 | 0.50 | 0.00 |
| Duck | 0.00 | 0.00 | 0.00 | 0.00 | 0.00 | 0.00 | 4.69 | 0.11 | 0.52 |

| | | | | | | | | | |
|---|---|---|---|---|---|---|---|---|---|
| Chicken | 0.00 | 0.00 | 0.00 | 0.00 | 0.00 | 0.00 | 1.33 | 0.44 | 0.59 |
| TOTAL (Average) | | | 0.65 | | | 0.09 | | | 0.46 |

**Supplementary Table 6.** Model of carbohydrate consumption over 9000 years in Asia, to construct Diet Model 1. The Total columns for each geographic region represents the proportion of carbohydrates contributed by the specific crop in each geographic region, respectively.

| CARBOHYDRATES | | | | | | | | | |
|---|---|---|---|---|---|---|---|---|---|
| | Southwest Asia | | | South Asia | | | East Asia | | |
| | Carb content | Proportion of years (out of 9000) of domestication | Total | Carb content | Proportion of years (out of 9000) of domestication | Total | Carb content | Proportion of years (out of 9000) of domestication | Total |
| Wheat | 42.50 | 1.00 | 42.50 | 0.00 | 0.00 | 0.00 | 0.00 | 0.00 | 0.00 |
| Barley | 77.70 | 1.00 | 77.70 | 0.00 | 0.00 | 0.00 | 0.00 | 0.00 | 0.00 |
| Lentil | 60.10 | 1.00 | 60.10 | 0.00 | 0.00 | 0.00 | 0.00 | 0.00 | 0.00 |
| Pea | 14.50 | 0.94 | 13.69 | 0.00 | 0.00 | 0.00 | 0.00 | 0.00 | 0.00 |
| Chickpea | 60.60 | 0.92 | 55.55 | 0.00 | 0.00 | 0.00 | 0.00 | 0.00 | 0.00 |
| Broadbean | 58.30 | 1.17 | 68.02 | 0.00 | 0.00 | 0.00 | 0.00 | 0.00 | 0.00 |
| Flax | 28.90 | 1.06 | 30.51 | 0.00 | 0.00 | 0.00 | 0.00 | 0.00 | 0.00 |
| Olive | 3.80 | 0.67 | 2.53 | 0.00 | 0.00 | 0.00 | 0.00 | 0.00 | 0.00 |
| Sheep | 0.00 | 0.89 | 0.00 | 0.00 | 0.00 | 0.00 | 0.00 | 0.00 | 0.00 |
| Goat | 0.00 | 0.89 | 0.00 | 0.00 | 0.00 | 0.00 | 0.00 | 0.00 | 0.00 |
| Cattle(taurine) | 0.00 | 0.89 | 0.00 | 0.00 | 0.00 | 0.00 | 0.00 | 0.00 | 0.00 |
| Cat | 0.00 | 0.44 | 0.00 | 0.00 | 0.00 | 0.00 | 0.00 | 0.00 | 0.00 |
| Tree Cotton | 0.00 | 0.00 | 0.00 | 0.00 | 0.50 | 0.00 | 0.00 | 0.00 | 0.00 |
| Rice (indica) | 0.00 | 0.00 | 0.00 | 74.90 | 0.28 | 20.80 | 0.00 | 0.00 | 0.00 |
| Little Millet | 0.00 | 0.00 | 0.00 | 72.90 | 0.50 | 36.45 | 0.00 | 0.00 | 0.00 |
| Browntop Millet | 0.00 | 0.00 | 0.00 | 72.90 | 0.44 | 32.40 | 0.00 | 0.00 | 0.00 |
| Mungbean | 0.00 | 0.00 | 0.00 | 62.60 | 0.33 | 20.87 | 0.00 | 0.00 | 0.00 |
| Pigeonpea | 0.00 | 0.00 | 0.00 | 23.90 | 0.39 | 9.29 | 0.00 | 0.00 | 0.00 |
| Cattle (zebu) | 0.00 | 0.00 | 0.00 | 0.00 | 0.72 | 0.00 | 0.00 | 0.00 | 0.00 |
| Water buffalo | 0.00 | 0.00 | 0.00 | 0.00 | 0.50 | 0.00 | 0.00 | 0.00 | 0.00 |
| Broomcorn millet | 0.00 | 0.00 | 0.00 | 0.00 | 0.00 | 0.00 | 72.90 | 0.89 | 64.80 |
| Foxtail millet | 0.00 | 0.00 | 0.00 | 0.00 | 0.00 | 0.00 | 72.90 | 0.83 | 60.75 |
| Rice (japonica) | 0.00 | 0.00 | 0.00 | 0.00 | 0.00 | 0.00 | 74.90 | 0.56 | 41.61 |
| Soybean | 0.00 | 0.00 | 0.00 | 0.00 | 0.00 | 0.00 | 30.20 | 0.39 | 11.74 |
| Ramie | 0.00 | 0.00 | 0.00 | 0.00 | 0.00 | 0.00 | 5.80 | 0.58 | 3.38 |
| Melon | 0.00 | 0.00 | 0.00 | 0.00 | 0.00 | 0.00 | 9.10 | 0.39 | 3.54 |
| Pig | 0.00 | 1.00 | 0.00 | 0.00 | 0.00 | 0.00 | 0.00 | 0.67 | 0.00 |

| | | | | | | | | | |
|---|---|---|---|---|---|---|---|---|---|
| Silkworm | 0.00 | 0.00 | 0.00 | 0.00 | 0.00 | 0.00 | 25.43 | 0.64 | 16.25 |
| Yak | 0.00 | 0.00 | 0.00 | 0.00 | 0.00 | 0.00 | 0.00 | 0.47 | 0.00 |
| Horse | 0.00 | 0.00 | 0.00 | 0.00 | 0.00 | 0.00 | 0.00 | 0.44 | 0.00 |
| Bactrian Camel | 0.00 | 0.00 | 0.00 | 0.00 | 0.00 | 0.00 | 4.60 | 0.50 | 2.30 |
| Duck | 0.00 | 0.00 | 0.00 | 0.00 | 0.00 | 0.00 | 0.00 | 0.11 | 0.00 |
| Chicken | 0.00 | 0.00 | 0.00 | 0.00 | 0.00 | 0.00 | 0.00 | 0.44 | 0.00 |
| TOTAL (Average) | | | 10.62 | | | 3.63 | | | 6.19 |

**Supplementary Table 7.** Model of omega-3 consumption over 9000 years in Asia, to construct Diet Model 1. The Total columns for each geographic region represents the proportion of omega-3 (g/100g) contributed by the specific crop in each geographic region, respectively.

| OMEGA 3 | | | | | | | | | |
|---|---|---|---|---|---|---|---|---|---|
| | Southwest Asia | | | South Asia | | | East Asia | | |
| | g/100g | Proportion of years (out of 9000) of domestication | Total | g/100g | Proportion of years (out of 9000) of domestication | Total | g/100g | Proportion of years (out of 9000) of domestication | Total |
| Wheat | 0.03 | 1.00 | 0.03 | 0.00 | 0.00 | 0.00 | 0.00 | 0.00 | 0.00 |
| Barley | 0.06 | 1.00 | 0.06 | 0.00 | 0.00 | 0.00 | 0.00 | 0.00 | 0.00 |
| Lentil | 0.11 | 1.00 | 0.11 | 0.00 | 0.00 | 0.00 | 0.00 | 0.00 | 0.00 |
| Pea | 0.04 | 0.94 | 0.03 | 0.00 | 0.00 | 0.00 | 0.00 | 0.00 | 0.00 |
| Chickpea | 0.10 | 0.92 | 0.09 | 0.00 | 0.00 | 0.00 | 0.00 | 0.00 | 0.00 |
| Broadbean | 0.05 | 1.17 | 0.05 | 0.00 | 0.00 | 0.00 | 0.00 | 0.00 | 0.00 |
| Flax | 22.81 | 1.06 | 24.08 | 0.00 | 0.00 | 0.00 | 0.00 | 0.00 | 0.00 |
| Olive | 0.09 | 0.67 | 0.06 | 0.00 | 0.00 | 0.00 | 0.00 | 0.00 | 0.00 |
| Sheep | 0.00 | 0.89 | 0.00 | 0.00 | 0.00 | 0.00 | 0.00 | 0.00 | 0.00 |
| Goat | 0.02 | 0.89 | 0.02 | 0.00 | 0.00 | 0.00 | 0.00 | 0.00 | 0.00 |
| Cattle/ (taurine) | 0.24 | 0.89 | 0.21 | 0.00 | 0.00 | 0.00 | 0.00 | 0.00 | 0.00 |
| Cat | 0.00 | 0.44 | 0.00 | 0.00 | 0.00 | 0.00 | 0.00 | 0.00 | 0.00 |
| Tree Cotton | 0.00 | 0.00 | 0.00 | 0.00 | 0.50 | 0.00 | 0.00 | 0.00 | 0.00 |
| Rice (indica) | 0.00 | 0.00 | 0.00 | 0.30 | 0.28 | 0.08 | 0.00 | 0.00 | 0.00 |
| Little Millet | 0.00 | 0.00 | 0.00 | 0.12 | 0.50 | 0.06 | 0.00 | 0.00 | 0.00 |
| Browntop Millet | 0.00 | 0.00 | 0.00 | 0.12 | 0.44 | 0.05 | 0.00 | 0.00 | 0.00 |
| Mungbean | 0.00 | 0.00 | 0.00 | 0.06 | 0.33 | 0.02 | 0.00 | 0.00 | 0.00 |
| Pigeonpea | 0.00 | 0.00 | 0.00 | 0.04 | 0.39 | 0.01 | 0.00 | 0.00 | 0.00 |
| Cattle (zebu) | 0.00 | 0.00 | 0.00 | 0.24 | 0.72 | 0.17 | 0.00 | 0.00 | 0.00 |
| Water buffalo | 0.00 | 0.00 | 0.00 | 0.04 | 0.50 | 0.02 | 0.00 | 0.00 | 0.00 |
| Broomcorn millet | 0.00 | 0.00 | 0.00 | 0.00 | 0.00 | 0.00 | 0.12 | 0.89 | 0.10 |

| | | | | | | | | | |
|---|---|---|---|---|---|---|---|---|---|
| Foxtail millet | 0.00 | 0.00 | 0.00 | 0.00 | 0.00 | 0.00 | 0.12 | 0.83 | 0.10 |
| Rice (japonica) | 0.00 | 0.00 | 0.00 | 0.00 | 0.00 | 0.00 | 0.30 | 0.56 | 0.17 |
| Soybean | 0.00 | 0.00 | 0.00 | 0.00 | 0.00 | 0.00 | 1.33 | 0.39 | 0.52 |
| Ramie | 0.00 | 0.00 | 0.00 | 0.00 | 0.00 | 0.00 | 0.00 | 0.58 | 0.00 |
| Melon | 0.00 | 0.00 | 0.00 | 0.00 | 0.00 | 0.00 | 0.03 | 0.39 | 0.01 |
| Pig | 0.00 | 1.00 | 0.00 | 0.00 | 0.00 | 0.00 | 0.43 | 0.67 | 0.28 |
| Silkworm | 0.00 | 0.00 | 0.00 | 0.00 | 0.00 | 0.00 | 0.00 | 0.64 | 0.00 |
| Yak | 0.00 | 0.00 | 0.00 | 0.00 | 0.00 | 0.00 | 0.00 | 0.47 | 0.00 |
| Horse | 0.00 | 0.00 | 0.00 | 0.00 | 0.00 | 0.00 | 0.36 | 0.44 | 0.16 |
| Bactrian Camel | 0.00 | 0.00 | 0.00 | 0.00 | 0.00 | 0.00 | 0.00 | 0.50 | 0.00 |
| Duck | 0.00 | 0.00 | 0.00 | 0.00 | 0.00 | 0.00 | 0.39 | 0.11 | 0.04 |
| Chicken | 0.00 | 0.00 | 0.00 | 0.00 | 0.00 | 0.00 | 0.10 | 0.44 | 0.04 |
| TOTAL (Average) | | | 0.75 | | | 0.01 | | | 0.04 |

**Supplementary Table 8.** Model of protein consumption over 9000 years in Asia, to construct Diet Model 1. The Total columns for each geographic region represents the proportion of protein (g/100g) contributed by the specific crop in each geographic region, respectively.

| PROTEIN | | | | | | | | | |
|---|---|---|---|---|---|---|---|---|---|
| | Southwest Asia | | | South Asia | | | East Asia | | |
| | g/100g | Proportion of years (out of 9000) of domestication | Total | g/100g | Proportion of years (out of 9000) of domestication | Total | g/100g | Proportion of years (out of 9000) of domestication | Total |
| Wheat | 7.50 | 1.00 | 7.50 | 0.00 | 0.00 | 0.00 | 0.00 | 0.00 | 0.00 |
| Barley | 2.30 | 1.00 | 2.30 | 0.00 | 0.00 | 0.00 | 0.00 | 0.00 | 0.00 |
| Lentil | 9.00 | 1.00 | 9.00 | 0.00 | 0.00 | 0.00 | 0.00 | 0.00 | 0.00 |
| Pea | 7.00 | 0.94 | 6.61 | 0.00 | 0.00 | 0.00 | 0.00 | 0.00 | 0.00 |
| Chickpea | 19.30 | 0.92 | 17.69 | 0.00 | 0.00 | 0.00 | 0.00 | 0.00 | 0.00 |
| Broadbean | 19.70 | 1.17 | 22.98 | 0.00 | 0.00 | 0.00 | 0.00 | 0.00 | 0.00 |
| Flax | 18.30 | 1.06 | 19.32 | 0.00 | 0.00 | 0.00 | 0.00 | 0.00 | 0.00 |
| Olive | 1.00 | 0.67 | 0.67 | 0.00 | 0.00 | 0.00 | 0.00 | 0.00 | 0.00 |
| Sheep | 20.60 | 0.89 | 18.31 | 0.00 | 0.00 | 0.00 | 0.00 | 0.00 | 0.00 |
| Goat | 27.10 | 0.89 | 24.09 | 0.00 | 0.00 | 0.00 | 0.00 | 0.00 | 0.00 |
| Cattle(taurine) | 26.00 | 0.89 | 23.11 | 0.00 | 0.00 | 0.00 | 0.00 | 0.00 | 0.00 |
| Cat | 0.00 | 0.44 | 0.00 | 0.00 | 0.00 | 0.00 | 0.00 | 0.00 | 0.00 |
| Tree Cotton | 0.00 | 0.00 | 0.00 | 0.00 | 0.50 | 0.00 | 0.00 | 0.00 | 0.00 |
| Rice (indica) | 0.00 | 0.00 | 0.00 | 4.00 | 0.28 | 1.11 | 0.00 | 0.00 | 0.00 |
| Little Millet | 0.00 | 0.00 | 0.00 | 3.50 | 0.50 | 1.75 | 0.00 | 0.00 | 0.00 |
| Browntop | 0.00 | 0.00 | 0.00 | 3.50 | 0.44 | 1.56 | 0.00 | 0.00 | 0.00 |

| | | | | | | | | | |
|---|---|---|---|---|---|---|---|---|---|
| Millet | | | | | | | | | |
| Mungbean | 0.00 | 0.00 | 0.00 | 7.00 | 0.33 | 2.33 | 0.00 | 0.00 | 0.00 |
| Pigeonpea | 0.00 | 0.00 | 0.00 | 6.00 | 0.39 | 2.33 | 0.00 | 0.00 | 0.00 |
| Cattle (zebu) | 0.00 | 0.00 | 0.00 | 26.00 | 0.72 | 18.78 | 0.00 | 0.00 | 0.00 |
| Water buffalo | 0.00 | 0.00 | 0.00 | 26.80 | 0.50 | 13.40 | 0.00 | 0.00 | 0.00 |
| Broomcorn millet | 0.00 | 0.00 | 0.00 | 0.00 | 0.00 | 0.00 | 3.50 | 0.89 | 3.11 |
| Foxtail millet | 0.00 | 0.00 | 0.00 | 0.00 | 0.00 | 0.00 | 3.50 | 0.44 | 1.56 |
| Rice (japonica) | 0.00 | 0.00 | 0.00 | 0.00 | 0.00 | 0.00 | 4.00 | 0.56 | 2.22 |
| Soybean | 0.00 | 0.00 | 0.00 | 0.00 | 0.00 | 0.00 | 16.60 | 0.39 | 6.46 |
| Ramie | 0.00 | 0.00 | 0.00 | 0.00 | 0.00 | 0.00 | 3.70 | 0.58 | 2.16 |
| Melon | 0.00 | 0.00 | 0.00 | 0.00 | 0.00 | 0.00 | 0.50 | 0.39 | 0.19 |
| Pig | 25.70 | 1.00 | 25.70 | 0.00 | 0.00 | 0.00 | 25.70 | 0.67 | 17.13 |
| Silkworm | 0.00 | 0.00 | 0.00 | 0.00 | 0.00 | 0.00 | 61.80 | 0.64 | 39.48 |
| Yak | 0.00 | 0.00 | 0.00 | 0.00 | 0.00 | 0.00 | 20.35 | 0.47 | 9.61 |
| Horse | 0.00 | 0.00 | 0.00 | 0.00 | 0.00 | 0.00 | 28.10 | 0.44 | 12.49 |
| Bactrian Camel | 0.00 | 0.00 | 0.00 | 0.00 | 0.00 | 0.00 | 2.80 | 0.50 | 1.40 |
| Duck | 0.00 | 0.00 | 0.00 | 0.00 | 0.00 | 0.00 | 19.00 | 0.11 | 2.11 |
| Chicken | 0.00 | 0.00 | 0.00 | 0.00 | 0.00 | 0.00 | 25.90 | 0.44 | 11.51 |
| TOTAL (Average) | | | 5.37 | | | 1.25 | | | 3.32 |

**Supplementary Table 9.** Dietary model 2, modeling glycemic index and carbohydrate content (g/100g) in each food item. The numbers next to each food item corresponds to the identification number of the item in the Diogenes GI database. Food items that do not have these numbers come from the Glycemic Index database.

| **Population** | **Population group** | **occupation** | 9000-0 YBP | | | | |
|---|---|---|---|---|---|---|---|
| India Sino-Tibetan | China, SE Asia, Sino-Tibet | hunter-gatherer | food item | % in diet | g carb/100g serving | carbs contributed by food item | GI |
| | | | acorns (Quercus emoryi), stewed with venison (F-P02, even carb) | 25 | 6.00 | 1.50 | 16.00 |
| | | | carp | 15 | 0.00 | 0.00 | 70.00 |
| | | | pork (UK-8055) | 10 | 0.40 | 0.04 | 70.00 |
| | | | peaches, raw (UK-944) | 10 | 7.60 | 0.76 | 42.00 |
| | | | melon.musk/cantaloupe, raw (UK-2533) | 10 | 5.00 | 0.50 | 72.00 |
| | | | raw vegetables salad (DE-1450) | 10 | 7.20 | 0.72 | 45.00 |

| | | | | | | | |
|---|---|---|---|---|---|---|---|
| | | | white yam (dioscorea alata), peeled, cubed, boiled (F-P02) | 15 | 20.67 | 3.10 | 75.00 |
| | | | oranges | 5 | 9.17 | 0.46 | 40.00 |
| | | | HUNTER-GATHERER 1 DIET | | | 7.08 | 53.40 |

| | | | | | | | |
|---|---|---|---|---|---|---|---|
| | | | 9000-8000 YBP | | | | |
| Tujia | China | agriculture | food item | % in diet | g carb/100g serving | carbs contributed by food item | GI |
| Miaozu | | | HUNTER-GATHERER 1 DIET | | | 7.08 | 53.40 |
| Han | | | | | | | |
| She | | | 8000-7000 YBP | | | | |
| | | | food item | % in diet | g carb/100g serving | carbs contributed by food item | GI |
| | | | japonica short-grain brown rice | 13 | 28.00 | 3.64 | 62.00 |
| | | | rice noodles, dried, boiled (Thai World, Bangkok, Thailand) | 3 | 21.67 | 0.65 | 61.00 |
| | | | rice porridge | 4 | 22.00 | 0.88 | 69.00 |
| | | | waxy (0-2% amylose) | 15 | 28.67 | 4.30 | 88.00 |
| | | | hunter-gatherer diet: | 15 | 7.08 | 1.06 | 53.40 |
| | | | hunter-gatherer diet: | 5 | 7.08 | 0.35 | 53.40 |
| | | | deer with acorns | 20 | 6.00 | 1.20 | 16.00 |
| | | | pork (UK-8055) | 5 | 0.40 | 0.02 | 70.00 |
| | | | peaches, raw (UK-944) | 10 | 7.60 | 0.76 | 45.00 |
| | | | melon.musk/cantaloupe, raw (UK-2533) | 10 | 5.00 | 0.50 | 72.00 |
| | | | 9000-8000 YBP DIET: | | | 13.37 | 65.16 |
| | | | 7000-0 YBP | | | | |
| | | | food item | % in diet | g carb/100g serving | carbs contributed by food item | GI |
| | | | japonica short-grain brown rice | 16 | 28.00 | 4.48 | 62.00 |
| | | | rice noodles, dried, boiled (Thai World, Bangkok, Thailand) | 3 | 21.67 | 0.65 | 61.00 |
| | | | rice porridge | 4 | 22.00 | 0.88 | 69.00 |
| | | | waxy (0-2% amylose) | 12 | 28.67 | 3.44 | 88.00 |
| | | | millet porridge | 15 | 24.00 | 3.60 | 62.00 |

| | | | | | | | |
|---|---|---|---|---|---|---|---|
| | | | lentils with vegetables, eaten with orange | 5 | 10.50 | 0.53 | 35.00 |
| | | | deer with acorns | 20 | 6.00 | 1.20 | 16.00 |
| | | | pork (UK-8055) | 5 | 0.40 | 0.02 | 70.00 |
| | | | peaches, raw (UK-944) | 10 | 7.60 | 0.76 | 42.00 |
| | | | melon.musk/cantaloupe, raw (UK-2533) | 10 | 5.00 | 0.50 | 72.00 |
| | | | 7000-0 YBP DIET: | | | 16.06 | 62.97 |
| | | | TOTAL DIET, since 9000 YBP | | | 14.76 | 62.15 |

| | | | | | | | |
|---|---|---|---|---|---|---|---|
| | | | 9000-8000 YBP | | | | |
| Cambodians | SE Asia | agriculture | food item | % in diet | g carb/100g serving | carbs contributed by food item | GI |
| | | | hunter-gatherer diet: | | | 7.08 | 53.40 |
| | | | 8000-6000 YBP | | | | |
| | | | food item | % in diet | g carb/100g serving | % carbs contributed by food item | GI |
| | | | japonica short-grain brown rice | 18 | 28.00 | 5.04 | 62.00 |
| | | | rice noodles, dried, boiled (Thai World, Bangkok, Thailand) | 3 | 21.67 | 0.65 | 61.00 |
| | | | rice porridge | 9 | 22.00 | 1.98 | 69.00 |
| | | | hunter-gatherer diet: | 10 | 7.08 | 0.71 | 53.40 |
| | | | raw vegetables salad (DE-1450) | 10 | 7.20 | 0.72 | 45.00 |
| | | | lentils with vegetables, eaten with orange | 5 | 10.50 | 0.53 | 35.00 |
| | | | deer with acorns | 20 | 6.00 | 1.20 | 16.00 |
| | | | pork (UK-8055) | 5 | 0.40 | 0.02 | 70.00 |
| | | | peaches, raw (UK-944) | 10 | 7.60 | 0.76 | 42.00 |
| | | | melon.musk/cantaloupe, raw (UK-2533) | 10 | 5.00 | 0.50 | 72.00 |
| | | | TOTAL: | | | 12.10 | 55.02 |
| | | | 6000-0 YBP | | | | |
| | | | food item | % in diet | g carb/100g serving | carbs contributed by food item | GI |
| | | | japonica short-grain brown rice | 18 | 28.00 | 5.04 | 62.00 |

| | | | food item | % in diet | g carb/100g serving | carbs contributed by food item | GI |
|---|---|---|---|---|---|---|---|
| | | | rice noodles, dried, boiled (Thai World, Bangkok, Thailand) | 3 | 21.67 | 0.65 | 61.00 |
| | | | rice porridge | 9 | 22.00 | 1.98 | 69.00 |
| | | | waxy (0-2% amylose) | 10 | 28.67 | 2.87 | 88.00 |
| | | | raw vegetables salad (DE-1450) | 10 | 7.20 | 0.72 | 45.00 |
| | | | lentils with vegetables, eaten with orange | 5 | 10.50 | 0.53 | 35.00 |
| | | | deer with acorns | 20 | 6.00 | 1.20 | 16.00 |
| | | | pork (UK-8055) | 5 | 0.40 | 0.02 | 70.00 |
| | | | peaches, raw (UK-944) | 10 | 7.60 | 0.76 | 42.00 |
| | | | melon.musk/cantaloupe, raw (UK-2533) | 10 | 5.00 | 0.50 | 72.00 |
| | | | TOTAL: | | | 14.26 | 61.73 |
| | | | TOTAL DIET, since 9000 YBP | | | 12.98 | 59.31 |

| | | | 9000-7000 YBP | | | | |
|---|---|---|---|---|---|---|---|
| Dai | SE Asia | agriculture | food item | % in diet | g carb/100g serving | carbs contributed by food item | GI |
| | | | hunter-gatherer diet: | | | 7.08 | 53.40 |
| | | | 7000-5500 YBP | | | | |
| | | | food item | % in diet | g carb/100g serving | carbs contributed by food item | GI |
| | | | hunter-gatherer diet: | 16 | 7.08 | 1.13 | 53.40 |
| | | | hunter-gatherer diet: | 3 | 7.08 | 0.21 | 53.40 |
| | | | hunter-gatherer diet: | 9 | 7.08 | 0.64 | 53.40 |
| | | | hunter-gatherer diet: | 12 | 7.08 | 0.85 | 53.40 |
| | | | millet porridge | 5 | 24.00 | 1.20 | 62.00 |
| | | | raw vegetables salad (DE-1450) | 5 | 7.20 | 0.36 | 45.00 |
| | | | lentils with vegetables, eaten with orange | 5 | 10.50 | 0.53 | 35.00 |
| | | | deer with acorns | 20 | 6.00 | 1.20 | 16.00 |
| | | | pork (UK-8055) | 5 | 0.40 | 0.02 | 70.00 |
| | | | peaches, raw (UK-944) | 10 | 7.60 | 0.76 | 42.00 |
| | | | melon.musk/cantaloupe, raw (UK-2533) | 10 | 5.00 | 0.50 | 72.00 |

| | | | | | | | |
|---|---|---|---|---|---|---|---|
| | | | TOTAL: | | | 7.40 | 47.14 |
| | | | 5500-0 YBP | | | | |
| | | | food item | % in diet | g carb/100g serving | carbs contributed by food item | GI |
| | | | japonica short-grain brown rice | 18 | 28.00 | 5.04 | 62.00 |
| | | | rice noodles, dried, boiled (Thai World, Bangkok, Thailand) | 3 | 21.67 | 0.65 | 61.00 |
| | | | rice porridge | 9 | 22.00 | 1.98 | 69.00 |
| | | | waxy (0-2% amylose) | 10 | 28.67 | 2.87 | 88.00 |
| | | | millet porridge | 5 | 24.00 | 1.20 | 62.00 |
| | | | raw vegetables salad (DE-1450) | 5 | 7.20 | 0.36 | 45.00 |
| | | | lentils with vegetables, eaten with orange | 5 | 10.50 | 0.53 | 35.00 |
| | | | deer with acorns | 20 | 6.00 | 1.20 | 16.00 |
| | | | pork (UK-8055) | 5 | 0.40 | 0.02 | 70.00 |
| | | | peaches, raw (UK-944) | 10 | 7.60 | 0.76 | 42.00 |
| | | | melon.musk/cantaloupe, raw (UK-2533) | 10 | 5.00 | 0.50 | 72.00 |
| | | | TOTAL: | | | 15.10 | 62.15 |
| | | | TOTAL DIET, since 9000 YBP | | | 12.03 | 56.70 |

| | | | | | | | |
|---|---|---|---|---|---|---|---|
| | | | 9000-4000 YBP | | | | |
| Burmese | SE Asia | agriculture | food item | % in diet | g carb/100g serving | carbs contributed by food item | GI |
| | | | hunter-gatherer diet: | | | 7.08 | 53.40 |
| | | | 4000-3000 YBP | | | | |
| | | | food item | % | g carb /100g serving | carbs contributed by food item | GI |
| | | | hunter-gatherer diet: | 16 | 7.08 | 1.13 | 53.40 |
| | | | hunter-gatherer diet: | 3 | 7.08 | 0.21 | 53.40 |
| | | | hunter-gatherer diet: | 9 | 7.08 | 0.64 | 53.40 |
| | | | hunter-gatherer diet: | 12 | 7.08 | 0.85 | 53.40 |
| | | | millet porridge | 5 | 24.00 | 1.20 | 62.00 |

| | | | | | | | |
|---|---|---|---|---|---|---|---|
| | | | raw vegetables salad (DE-1450) | 5 | 7.20 | 0.36 | 45.00 |
| | | | lentils with vegetables, eaten with orange | 5 | 10.50 | 0.53 | 35.00 |
| | | | deer with acorns | 20 | 6.00 | 1.20 | 16.00 |
| | | | pork (UK-8055) | 5 | 0.40 | 0.02 | 70.00 |
| | | | peaches, raw (UK-944) | 10 | 7.60 | 0.76 | 42.00 |
| | | | melon.musk/cantaloupe, raw (UK-2533) | 10 | 5.00 | 0.50 | 72.00 |
| | | | TOTAL AGRIC DIET: | | | 7.40 | 47.14 |
| | | | 3000-0 YBP | | | | |
| | | | food item | % | g carb /100g serving | carbs contributed by food item | GI |
| | | | japonica short-grain brown rice | 16 | 28.00 | 4.48 | 62.00 |
| | | | rice noodles, dried, boiled (Thai World, Bangkok, Thailand) | 3 | 21.67 | 0.65 | 61.00 |
| | | | rice porridge | 9 | 22.00 | 1.98 | 69.00 |
| | | | waxy (0-2% amylose) | 12 | 28.67 | 3.44 | 88.00 |
| | | | millet porridge | 5 | 24.00 | 1.20 | 62.00 |
| | | | raw vegetables salad (DE-1450) | 5 | 7.20 | 0.36 | 45.00 |
| | | | lentils with vegetables, eaten with orange | 5 | 10.50 | 0.53 | 35.00 |
| | | | deer with acorns | 20 | 6.00 | 1.20 | 16.00 |
| | | | pork (UK-8055) | 5 | 0.40 | 0.02 | 70.00 |
| | | | peaches, raw (UK-944) | 10 | 7.60 | 0.76 | 42.00 |
| | | | melon.musk/cantaloupe, raw (UK-2533) | 10 | 5.00 | 0.50 | 72.00 |
| | | | TOTAL AGRIC DIET: | | | 15.12 | 63.13 |
| | | | TOTAL DIET, since 9000 YBP | | | 9.79 | 55.95 |

| | | | | | | | |
|---|---|---|---|---|---|---|---|
| | | | 9000-7000 YBP | | | | |
| Japanese | East Asia | agriculture | food item | % in diet | g carb/100g serving | carbs contributed by food item | GI |
| | | | hunter-gatherer diet: | | | 7.08 | 53.40 |
| | | | 7000-3000 YBP | | | | |

| food item | % | g carb /100g serving | carbs contributed by food item | GI |
|---|---|---|---|---|
| hunter-gatherer diet: | 16 | 7.08 | 1.13 | 53.40 |
| hunter-gatherer diet: | 3 | 7.08 | 0.21 | 53.40 |
| hunter-gatherer diet: | 4 | 7.08 | 0.28 | 53.40 |
| hunter-gatherer diet: | 12 | 7.08 | 0.85 | 53.40 |
| millet porridge | 15 | 24.00 | 3.60 | 62.00 |
| lentils with vegetables, eaten with orange | 5 | 10.50 | 0.53 | 35.00 |
| deer with acorns | 20 | 6.00 | 1.20 | 16.00 |
| pork (UK-8055) | 5 | 0.40 | 0.02 | 70.00 |
| peaches, raw (UK-944) | 10 | 7.60 | 0.76 | 42.00 |
| melon.musk/cantaloupe, raw (UK-2533) | 10 | 5.00 | 0.50 | 72.00 |
| TOTAL AGRIC DIET: | | | 9.08 | 50.91 |

3000-0 YBP

| food item | % | g carb /100g serving | carbs contributed by food item | GI |
|---|---|---|---|---|
| japonica short-grain brown rice | 16 | 28.00 | 4.48 | 62.00 |
| rice noodles, dried, boiled (Thai World, Bangkok, Thailand) | 3 | 21.67 | 0.65 | 61.00 |
| rice porridge | 4 | 22.00 | 0.88 | 69.00 |
| waxy (0-2% amylose) | 12 | 28.67 | 3.44 | 88.00 |
| millet porridge | 15 | 24.00 | 3.60 | 62.00 |
| lentils with vegetables, eaten with orange | 5 | 10.50 | 0.53 | 35.00 |
| deer with acorns | 20 | 6.00 | 1.20 | 16.00 |
| pork (UK-8055) | 5 | 0.40 | 0.02 | 70.00 |
| peaches, raw (UK-944) | 10 | 7.60 | 0.76 | 42.00 |
| melon.musk/cantaloupe, raw (UK-2533) | 10 | 5.00 | 0.50 | 72.00 |
| TOTAL AGRIC DIET: | | | 16.06 | 62.97 |
| TOTAL DIET, since 9000 YBP | | | 10.96 | 55.48 |

9000-7000 YBP

| | | | | | | | |
|---|---|---|---|---|---|---|---|
| Tu | China | pastoral | food item | % in diet | g carb/100g serving | carbs contributed by food item | GI |
| | (calculating agricultural values) | | hunter-gatherer diet: | | | 7.08 | 53.40 |
| | | estimates for agricultural diet | | | | | |
| | | | 7000-5500 YBP | | | | |
| | | | food item | % | g carb /100g serving | carbs contributed by food item | GI |
| | | | hunter-gatherer diet: | 18 | 7.08 | 1.27 | 53.40 |
| | | | hunter-gatherer diet: | 3 | 7.08 | 0.21 | 53.40 |
| | | | hunter-gatherer diet: | 4 | 7.08 | 0.28 | 53.40 |
| | | | hunter-gatherer diet: | 10 | 7.08 | 0.71 | 53.40 |
| | | | millet porridge | 15 | 24.00 | 3.60 | 62.00 |
| | | | lentils with vegetables, eaten with orange | 5 | 10.50 | 0.53 | 35.00 |
| | | | deer with acorns | 20 | 6.00 | 1.20 | 16.00 |
| | | | pork (UK-8055) | 5 | 0.40 | 0.02 | 70.00 |
| | | | peaches, raw (UK-944) | 10 | 7.60 | 0.76 | 42.00 |
| | | | melon.musk/cantaloupe, raw (UK-2533) | 10 | 5.00 | 0.50 | 72.00 |
| | | | TOTAL AGRIC DIET: | | | 9.26 | 49.92 |
| | | | 5500-0 YBP | | | | |
| | | | food item | % | g carb /100g serving | carbs contributed by food item | GI |
| | | | japonica short-grain brown rice | 18 | 28.00 | 5.04 | 62.00 |
| | | | rice noodles, dried, boiled (Thai World, Bangkok, Thailand) | 3 | 21.67 | 0.65 | 61.00 |
| | | | rice porridge | 4 | 22.00 | 0.88 | 69.00 |
| | | | waxy (0-2% amylose) | 10 | 28.67 | 2.87 | 88.00 |
| | | | millet porridge | 15 | 24.00 | 3.60 | 62.00 |
| | | | lentils with vegetables, eaten with orange | 5 | 10.50 | 0.53 | 35.00 |
| | | | deer with acorns | 20 | 6.00 | 1.20 | 16.00 |
| | | | pork (UK-8055) | 5 | 0.40 | 0.02 | 70.00 |
| | | | peaches, raw (UK-944) | 10 | 7.60 | 0.76 | 42.00 |
| | | | melon.musk/cantaloupe, raw (UK-2533) | 10 | 5.00 | 0.50 | 72.00 |

| | | | | | | | |
|---|---|---|---|---|---|---|---|
| | | | TOTAL AGRIC DIET: | | | 16.04 | 62.04 |
| | | | EST. AGRIC DIET, since 9000 YBP | | | 12.92 | 58.10 |
| | | estimates for pastoral diet | 9000-5500 YBP | | | | |
| | | | food item | % in diet | g carb/100g serving | carbs contributed by food item | GI |
| | | | hunter-gatherer diet: | | | 7.08 | 53.40 |
| | | | 5500-4000 YBP | | | | |
| | | | food item | % | g carb /100g serving | carbs contributed by food item | GI |
| | | | hunter-gatherer diet: | 15 | 7.08 | 1.06 | 53.40 |
| | | | hunter-gatherer diet: | 15 | 7.08 | 1.06 | 53.40 |
| | | | hunter-gatherer diet: | 20 | 7.08 | 1.42 | 53.40 |
| | | | | 50 | 16.04 | 8.02 | 62.04 |
| | | | TOTAL PASTORAL DIET: | | | 11.56 | 59.39 |
| | | | 4000-0 YBP | | | | |
| | | | food item | % | g carb /100g serving | carbs contributed by food item | GI |
| | | | beef | 15 | 0.00 | 0.00 | 70.00 |
| | | | lamb | 15 | 0.00 | 0.00 | 70.00 |
| | | | whole milk (UK-277) | 20 | 4.50 | 0.90 | 34.00 |
| | | | Tu agricultural diet | 50 | 16.04 | 8.02 | 62.04 |
| | | | TOTAL PASTORAL DIET: | | | 8.92 | 59.21 |
| | | | PASTORAL DIET, since 9000 YBP | | | 8.65 | 56.82 |

| | | | | | | | |
|---|---|---|---|---|---|---|---|
| | | | 9000-6000 YBP | | | | |
| Yizu | China, Sino-Tibet | 0.66 pastoral, 0.33 agric | food item | % in diet | g carb/100g serving | carbs contributed by food item | GI |

| | | | | | |
|---|---|---|---|---|---|
| (calculating agricultural values) | hunter-gatherer diet: | | | 7.08 | 53.40 |

6000-4500 YBP

| | % | g carb /100g serving | carbs contributed by food item | GI |
|---|---|---|---|---|
| japonica short-grain brown rice | 18 | 7.08 | 1.27 | 53.40 |
| rice noodles, dried, boiled (Thai World, Bangkok, Thailand) | 3 | 7.08 | 0.21 | 53.40 |
| rice porridge | 4 | 7.08 | 0.28 | 53.40 |
| waxy (0-2% amylose) | 10 | 7.08 | 0.71 | 53.40 |
| millet porridge | 15 | 24.00 | 3.60 | 62.00 |
| lentils with vegetables, eaten with orange | 5 | 10.50 | 0.53 | 35.00 |
| deer with acorns | 20 | 6.00 | 1.20 | 16.00 |
| pork (UK-8055) | 5 | 0.40 | 0.02 | 70.00 |
| peaches, raw (UK-944) | 10 | 7.60 | 0.76 | 42.00 |
| melon.musk/cantaloupe, raw (UK-2533) | 10 | 5.00 | 0.50 | 72.00 |
| TOTAL AGRIC DIET: | | | 9.08 | 50.91 |

4500-0 YBP

| food item | % in diet | g carb/100g serving | carbs contributed by food item | GI |
|---|---|---|---|---|
| japonica short-grain brown rice | 18 | 28.00 | 5.04 | 62.00 |
| rice noodles, dried, boiled (Thai World, Bangkok, Thailand) | 3 | 21.67 | 0.65 | 61.00 |
| rice porridge | 4 | 22.00 | 0.88 | 69.00 |
| waxy (0-2% amylose) | 10 | 28.67 | 2.87 | 88.00 |
| millet porridge | 15 | 24.00 | 3.60 | 62.00 |
| lentils with vegetables, eaten with orange | 5 | 10.50 | 0.53 | 35.00 |
| deer with acorns | 20 | 6.00 | 1.20 | 16.00 |
| pork (UK-8055) | 5 | 0.40 | 0.02 | 70.00 |
| peaches, raw (UK-944) | 10 | 7.60 | 0.76 | 42.00 |
| melon.musk/cantaloupe, raw (UK-2533) | 10 | 5.00 | 0.50 | 72.00 |
| TOTAL AGRIC DIET: | | | 16.04 | 62.04 |

| | | | | | | | |
|---|---|---|---|---|---|---|---|
| | | | AGRIC DIET, since 9000 YBP | | | 11.89 | 57.30 |
| | | | 9000-4500 YBP | | | | |
| | | (calculating pastoral values) | food item | % in diet | g carb/100g serving | carbs contributed by food item | GI |
| | | | (H-G diet) | | | 7.08 | 53.40 |
| | | | 4500-4000 YBP | | | | |
| | | | food item | % in diet | g carb/100g serving | carbs contributed by food item | GI |
| | | | hunter-gatherer diet: | 15 | 7.08 | 1.06 | 53.40 |
| | | | hunter-gatherer diet: | 15 | 7.08 | 1.06 | 53.40 |
| | | | hunter-gatherer diet: | 20 | 7.08 | 1.42 | 53.40 |
| | | | | 50 | 16.04 | 8.02 | 62.04 |
| | | | | | | 11.56 | 59.39 |
| | | | 4000-0 YBP | | | | |
| | | | food item | % in diet | g carb/100g serving | carbs contributed by food item | GI |
| | | | beef | 15 | 0.00 | 0.00 | 70.00 |
| | | | lamb | 15 | 0.00 | 0.00 | 70.00 |
| | | | whole milk (UK-277) | 20 | 4.50 | 0.90 | 34.00 |
| | | | Yizu agricultural diet | 50 | 16.04 | 8.02 | 62.04 |
| | | | TOTAL PASTORAL DIET: | | | 8.92 | 59.21 |
| | | | PASTORAL DIET, since 9000 YBP | | | 8.15 | 59.61 |
| | | | TOTAL DIET, since 9000 YBP | | | 10.64 | 58.07 |

| | | | | | | | |
|---|---|---|---|---|---|---|---|
| | | | 9000-5000 YBP | | | | |
| Naxi | China, Sino-Tibet | 0.66 agric, 0.33 pastoral | food item | % in diet | g carb/100g serving | carbs contributed by food item | GI |
| | (calculating agricultural | | hunter-gatherer diet: | | | 7.08 | 53.40 |

values)

5000-3500 YBP

| food item | % in diet | g carb /100g serving | carbs contributed by food item | GI |
|---|---|---|---|---|
| japonica short-grain brown rice | 18 | 7.08 | 1.27 | 53.40 |
| rice noodles, dried, boiled (Thai World, Bangkok, Thailand) | 3 | 7.08 | 0.21 | 53.40 |
| rice porridge | 4 | 7.08 | 0.28 | 53.40 |
| waxy (0-2% amylose) | 10 | 7.08 | 0.71 | 53.40 |
| millet porridge | 15 | 24.00 | 3.60 | 62.00 |
| lentils with vegetables, eaten with orange | 5 | 10.50 | 0.53 | 35.00 |
| deer with acorns | 20 | 6.00 | 1.20 | 16.00 |
| pork (UK-8055) | 5 | 0.40 | 0.02 | 70.00 |
| peaches, raw (UK-944) | 10 | 7.60 | 0.76 | 42.00 |
| melon.musk/cantaloupe, raw (UK-2533) | 10 | 5.00 | 0.50 | 72.00 |
| TOTAL AGRIC DIET: | | | 9.08 | 50.91 |

3500-0 YBP

| food item | % in diet | g carb/100g serving | carbs contributed by food item | GI |
|---|---|---|---|---|
| japonica short-grain brown rice | 18 | 28.00 | 5.04 | 62.00 |
| rice noodles, dried, boiled (Thai World, Bangkok, Thailand) | 3 | 21.67 | 0.65 | 61.00 |
| rice porridge | 4 | 22.00 | 0.88 | 69.00 |
| waxy (0-2% amylose) | 10 | 28.67 | 2.87 | 88.00 |
| millet porridge | 15 | 24.00 | 3.60 | 62.00 |
| lentils with vegetables, eaten with orange | 5 | 10.50 | 0.53 | 35.00 |
| deer with acorns | 20 | 6.00 | 1.20 | 16.00 |
| pork (UK-8055) | 5 | 0.40 | 0.02 | 70.00 |
| peaches, raw (UK-944) | 10 | 7.60 | 0.76 | 42.00 |
| melon.musk/cantaloupe, raw (UK-2533) | 10 | 5.00 | 0.50 | 72.00 |
| TOTAL AGRIC DIET: | | | 16.04 | 62.04 |

| | | | | | | | |
|---|---|---|---|---|---|---|---|
| | | | AGRIC DIET, since 9000 YBP | | | 10.90 | 56.34 |
| | | | 9000-4000 YBP | | | | |
| | (calculating pastoral values) | | food item | | g carb /100g serving | carbs contributed by food item | GI |
| | | | hunter-gatherer diet: | | | 7.08 | 53.40 |
| | | | 4000-3500 YBP | | | | |
| | | | food item | % in diet | g carb /100g serving | carbs contributed by food item | GI |
| | | | hunter-gatherer diet: | 15 | 7.08 | 1.06 | 53.40 |
| | | | hunter-gatherer diet: | 15 | 7.08 | 1.06 | 53.40 |
| | | | hunter-gatherer diet: | 20 | 7.08 | 1.42 | 53.40 |
| | | | China agricultural diet | 50 | 16.04 | 8.02 | 62.04 |
| | | | TOTAL PASTORAL DIET: | | | 11.56 | 59.39 |
| | | | 3500-0 YBP | | | | |
| | | | food item | % in diet | g carb /100g serving | carbs contributed by food item | GI |
| | | | beef | 15 | 0.00 | 0.00 | 70.00 |
| | | | lamb | 15 | 0.00 | 0.00 | 70.00 |
| | | | whole milk (UK-277) | 20 | 4.50 | 0.90 | 34.00 |
| | | | China agricultural diet | 50 | 16.04 | 8.02 | 62.04 |
| | | | TOTAL PASTORAL DIET: | | | 8.92 | 59.21 |
| | | | PASTORAL DIET, since 9000 YBP | | | 9.33 | 62.59 |
| | | | TOTAL DIET, since 9000 YBP | | | 10.37 | 58.43 |

| | | | | | | | |
|---|---|---|---|---|---|---|---|
| | | | 9000-4500 YBP | | | | |
| Uygur | Xinjiang province, China | agricultural | food item | % in diet | g carb /100g serving | carbs contributed by food item | GI |
| | | | apple, raw, golden delicious | 30 | 13.33 | 4.00 | 39.00 |
| | | | apricot raw (DE-135) | 30 | 7.50 | 2.25 | 57.00 |

| | | | | |
|---|---|---|---|---|
| hunted meat (misc) | 40 | 0.00 | 0.00 | 70.00 |
| TOTAL UYGUR PRE-4500BP DIET: | | | 6.25 | 45.48 |

4500-4000 YBP

| food item | % in diet | g carb /100g serving | carbs contributed by food item | GI |
|---|---|---|---|---|
| steamed triticum aestivum | 15 | 74.63 | 11.19 | 55.00 |
| barley (hordeum vulgare) | 15 | 24.67 | 3.70 | 37.00 |
| TOTAL UYGUR PRE-4500BP DIET: | 15 | 6.25 | 0.94 | 45.48 |
| TOTAL UYGUR PRE-4500BP DIET: | 10 | 6.25 | 0.62 | 45.48 |
| melon.musk/cantaloupe, raw (UK-2533) | 10 | 5.00 | 0.50 | 72.00 |
| whole milk (UK-277) | 15 | 4.50 | 0.68 | 34.00 |
| apple, raw, golden delicious | 10 | 13.33 | 1.33 | 39.00 |
| apricot raw (DE-135) | 10 | 7.50 | 0.75 | 57.00 |
| TOTAL UYGUR DIET: | | | 19.72 | 47.14 |

4000-0 YBP

| food item | % in diet | g carb /100g serving | carbs contributed by food item | GI |
|---|---|---|---|---|
| steamed triticum aestivum | 15 | 74.63 | 11.19 | 55.00 |
| barley (hordeum vulgare) | 15 | 24.67 | 3.70 | 37.00 |
| beef | 15 | 0.00 | 0.00 | 70.00 |
| lamb | 10 | 0.00 | 0.00 | 70.00 |
| melon.musk/cantaloupe, raw (UK-2533) | 10 | 5.00 | 0.50 | 72.00 |
| whole milk (UK-277) | 15 | 4.50 | 0.68 | 34.00 |
| apple, raw, golden delicious | 10 | 13.33 | 1.33 | 39.00 |
| apricot raw (DE-135) | 10 | 7.50 | 0.75 | 57.00 |
| TOTAL UYGUR DIET: | | | 18.15 | 47.28 |
| TOTAL DIET, since 9000 YBP | | | 12.29 | 46.37 |

| | | | | | | | |
|---|---|---|---|---|---|---|---|
| Xibo | Xinjiang province, China | part pastoral | 9000-4500 YBP | | | | |
| | rest: Uygur pre-4500BP | | food item | % in diet | g carb /100g serving | carbs contributed by food item | GI |
| | | | apple, raw, golden delicious | 30 | 13.33 | 4.00 | 39.00 |
| | | | apricot raw (DE-135) | 30 | 7.50 | 2.25 | 57.00 |
| | | | hunted meat (misc) | 40 | 0.00 | 0.00 | 70.00 |
| | | | TOTAL UYGUR PRE-4500BP DIET: | | | 6.25 | 45.48 |
| | | | 4500-4000 YBP | | | | |
| | | | food item | % in diet | g carb /100g serving | carbs contributed by food item | GI |
| | | | TOTAL UYGUR PRE-4500BP DIET: | 10 | 6.25 | 0.67 | 45.48 |
| | | | TOTAL UYGUR PRE-4500BP DIET: | 20 | 6.25 | 1.34 | 45.48 |
| | | | Uygur diet | 70 | 12.29 | 8.60 | 46.37 |
| | | | TOTAL XIBO DIET: | | | 10.48 | 46.21 |
| | | | 4000-0 YBP | | | | |
| | | | food item | % in diet | g carb /100g serving | carbs contributed by food item | GI |
| | | | whole milk (UK-277) | 10 | 5.00 | 0.50 | 34.00 |
| | | | lamb | 20 | 0.00 | 0.00 | 70.00 |
| | | | Uygur diet | 70 | 12.29 | 8.60 | 46.37 |
| | | | TOTAL XIBO DIET: | | | 9.10 | 45.69 |
| | | | TOTAL DIET, since 9000 YBP | | | 7.75 | 45.62 |

| | | | | | | | |
|---|---|---|---|---|---|---|---|
| | | | 9000-0 YBP | | | | |
| Central India tribe | India | hunter-gatherer | food item | % in diet | g carb /100g serving | carbs contributed by food item | GI |
| South India tribe | | | chicken and salad vegetables | 35 | 0.00 | 0.00 | 70.00 |
| | | | carrots, raw (UK-2405) | 20 | 5.40 | 1.08 | 47.00 |
| | | | white yam (dioscorea alata), peeled, cubed, boiled | 15 | 20.67 | 3.10 | 75.00 |

| | | | | | | | |
|---|---|---|---|---|---|---|---|
| | | | oranges | 10 | 9.17 | 0.92 | 40.00 |
| | | | deer | 10 | 0.00 | 0.00 | 70.00 |
| | | | Mango (Mangifera indica), raw | 10 | 12.50 | 1.25 | 51.00 |
| | | | TOTAL H-G DIET: | | | 6.35 | 60.45 |

| | | | | | | | |
|---|---|---|---|---|---|---|---|
| | | | 9000-5000 YBP | | | | |
| South India caste | South India | agriculture | food item | % in diet | g carb /100g serving | carbs contributed by food item | GI |
| | | | hunter-gatherer diet: | | | 6.35 | 60.45 |
| | | | 5000-4000 YBP | | | | |
| | | | food item | % | g carb /100g serving | % carbs contributed by food item | GI |
| | | | hunter-gatherer diet: | 18 | 6.35 | 1.14 | 60.45 |
| | | | hunter-gatherer diet: | 5 | 6.35 | 0.32 | 60.45 |
| | | | hunter-gatherer diet: | 10 | 6.35 | 0.63 | 60.45 |
| | | | hunter-gatherer diet: | 10 | 6.35 | 0.63 | 60.45 |
| | | | hunter-gatherer diet: | 7 | 6.35 | 0.44 | 60.45 |
| | | | hunter-gatherer diet: | 5 | 6.35 | 0.32 | 60.45 |
| | | | hunter-gatherer diet: | 5 | 6.35 | 0.32 | 60.45 |
| | | | beef (and other meat) | 20 | 0.00 | 0.00 | 70.00 |
| | | | oranges | 10 | 9.17 | 0.92 | 40.00 |
| | | | raw vegetables salad (DE-1450) | 10 | 7.20 | 0.72 | 45.00 |
| | | | TOTAL AGRICULTURAL DIET: | | | 5.45 | 54.97 |
| | | | 4000-3000 YBP | | | | |
| | | | food item | % | g carb /100g serving | carbs contributed by food item | GI |
| | | | (5000-4000 YBP diet) | 18 | 6.35 | 1.14 | 60.45 |
| | | | (5000-4000 YBP diet) | 5 | 6.35 | 0.32 | 60.45 |
| | | | (5000-4000 YBP diet) | 10 | 6.35 | 0.63 | 60.45 |
| | | | (5000-4000 YBP diet) | 10 | 6.35 | 0.63 | 60.45 |
| | | | laddu (popped amaranth, foxtail, roasted legume, fenugreek) in syrup | 7 | 62.00 | 4.34 | 24.00 |

| | | | | |
|---|---|---|---|---|
| black gram (phaseolus mungo), soaked, stored, steamed | 5 | 12.00 | 0.60 | 43.00 |
| horse gram (Dolichos biflorus), soaked, stored, steamed | 5 | 19.33 | 0.97 | 51.00 |
| beef (and other meat) | 20 | 0.00 | 0.00 | 70.00 |
| oranges | 10 | 9.17 | 0.92 | 40.00 |
| raw vegetables salad (DE-1450) | 10 | 7.20 | 0.72 | 45.00 |
| TOTAL AGRICULTURE DIET: | | | 10.27 | 40.24 |

3000-0 YBP

| food item | % | g carb /100g serving | carbs contributed by food item | GI |
|---|---|---|---|---|
| brown, boiled (South India) | 18 | 22.00 | 3.96 | 50.00 |
| Moolgiri white jasmine rice (Tajmahal Agro Industries) | 20 | 21.33 | 4.27 | 54.00 |
| roasted bread from jowar flour (Sorghum vulgare) | 10 | 71.43 | 7.14 | 77.00 |
| porridge, made from decorticated finger millet, eaten with Bengal gram, green gram, and black gram | 10 | 67.00 | 6.70 | 65.00 |
| laddu (popped amaranth, foxtail, roasted legume, fenugreek) in syrup | 7 | 62.00 | 4.34 | 24.00 |
| black gram (phaseolus mungo), soaked, stored, steamed | 5 | 12.00 | 0.60 | 43.00 |
| horse gram (Dolichos biflorus), soaked, stored, steamed | 5 | 19.33 | 0.97 | 51.00 |
| beef (and other meat) | 20 | 0.00 | 0.00 | 70.00 |
| oranges | 10 | 0.92 | 9.17 | 40.00 |
| raw vegetables salad (DE-1450) | 10 | 7.20 | 0.72 | 45.00 |
| TOTAL AGRICULTURE DIET: | | | 37.86 | 52.61 |
| TOTAL DIET, since 9000 YBP | | | 17.19 | 54.98 |

9000-5000 YBP

| North India caste | North India | agriculture | food item | % in diet | g carb /100g serving | carbs contributed by food item | GI |
|---|---|---|---|---|---|---|---|
| | | | hunter-gatherer diet: | | | 6.35 | 60.45 |

5000-4500 YBP

| food item | % in diet | g carb /100g serving | carbs contributed by food item | GI |
|---|---|---|---|---|
| (H-G diet) | 20 | 6.35 | 1.27 | 60.45 |
| (H-G diet) | 5 | 6.35 | 0.32 | 60.45 |
| (H-G diet) | 4 | 6.35 | 0.25 | 60.45 |
| (H-G diet) | 6 | 6.35 | 0.38 | 60.45 |
| (H-G diet) | 10 | 6.35 | 0.63 | 60.45 |
| lentils and vegetables, eaten with orange | 15 | 10.50 | 1.58 | 35.00 |
| deer | 10 | 0.00 | 0.00 | 70.00 |
| Brown and Wild rice, Uncle Ben's Ready Whole Grain Medley (pouch) | 10 | 26.00 | 2.60 | 45.00 |
| chicken | 10 | 0.00 | 0.00 | 70.00 |
| raw vegetables salad (DE-1450) | 10 | 7.20 | 0.72 | 45.00 |
| TOTAL AGRICULTURE DIET: | | | 7.75 | 48.66 |

4500-3000 YBP

| food item | % in diet | g carb /100g serving | carbs contributed by food item | GI |
|---|---|---|---|---|
| Moolgiri white jasmine rice (Tajmahal Agro Industries) | 20 | 21.33 | 4.27 | 54.00 |
| steamed triticum aestivum | 5 | 74.63 | 3.73 | 55.00 |
| barley (hordeum vulgare) | 4 | 24.67 | 0.99 | 48.00 |
| coarse barley kernel bread (80% scalded kernels, 20% white wheat flour) | 6 | 66.67 | 4.00 | 34.00 |
| beef (or other domesticated meat) | 10 | 0.00 | 0.00 | 70.00 |
| lentils and vegetables, eaten with orange | 15 | 10.50 | 1.58 | 35.00 |
| deer | 10 | 0.00 | 0.00 | 70.00 |

| | | | | | | | |
|---|---|---|---|---|---|---|---|
| | | | Brown and Wild rice, Uncle Ben's Ready Whole Grain Medley (pouch) | 10 | 26.00 | 2.60 | 45.00 |
| | | | chicken | 10 | 0.00 | 0.00 | 70.00 |
| | | | raw vegetables salad (DE-1450) | 10 | 7.20 | 0.72 | 45.00 |
| | | | TOTAL AGRICULTURE DIET: | | | 17.88 | 46.06 |
| | | | 3000-0 YBP | | | | |
| | | | food item | % in diet | g carb /100g serving | carbs contributed by food item | GI |
| | | | Moolgiri white jasmine rice (Tajmahal Agro Industries) | 25 | 21.33 | 5.33 | 54.00 |
| | | | steamed triticum aestivum | 6 | 74.63 | 4.66 | 55.00 |
| | | | barley (hordeum vulgare) | 5 | 24.67 | 1.23 | 48.00 |
| | | | coarse barley kernel bread (80% scalded kernels, 20% white wheat flour) | 8 | 66.67 | 5.00 | 34.00 |
| | | | beef (or other domesticated meat) | 13 | 0.00 | 0.00 | 70.00 |
| | | | lentils and vegetables, eaten with orange | 19 | 10.50 | 1.97 | 35.00 |
| | | | chicken | 13 | 0.00 | 0.00 | 70.00 |
| | | | raw vegetables salad (DE-1450) | 13 | 7.20 | 0.72 | 45.00 |
| | | | TOTAL AGRICULTURE DIET: | | | 18.92 | 46.25 |
| | | | TOTAL DIET, since 9000 YBP | | | 12.54 | 52.67 |

| | | | | | | | |
|---|---|---|---|---|---|---|---|
| | | | 9000-5000 YBP | | | | |
| Gujuratis | North India | agriculture | food item | % in diet | g carb /100g serving | carbs contributed by food item | GI |
| | | | hunter-gatherer diet: | | | 6.35 | 60.45 |
| | | | 5000-4500 YBP | | | | |
| | | | food item | % | g carb /100g serving | carbs contributed by food item | GI |
| | | | (H-G diet) | 20 | 6.35 | 1.27 | 60.45 |

| | | | | |
|---|---|---|---|---|
| (H-G diet) | 5 | 6.35 | 0.32 | 60.45 |
| (H-G diet) | 4 | 6.35 | 0.25 | 60.45 |
| (H-G diet) | 6 | 6.35 | 0.38 | 60.45 |
| (H-G diet) | 10 | 6.35 | 0.63 | 60.45 |
| lentils and vegetables, eaten with orange | 15 | 10.50 | 1.58 | 35.00 |
| deer | 10 | 0.00 | 0.00 | 70.00 |
| Brown and Wild rice, Uncle Ben's Ready Whole Grain Medley (pouch) | 10 | 26.00 | 2.60 | 45.00 |
| chicken | 10 | 0.00 | 0.00 | 70.00 |
| raw vegetables salad (DE-1450) | 10 | 7.20 | 0.72 | 45.00 |
| TOTAL AGRICULTURE DIET: | | | 7.75 | 48.66 |

4500-3000 YBP

| food item | % in diet | g carb /100g serving | carbs contributed by food item | GI |
|---|---|---|---|---|
| Moolgiri white jasmine rice (Tajmahal Agro Industries) | 20 | 21.33 | 4.27 | 54.00 |
| steamed triticum aestivum | 5 | 74.63 | 3.73 | 55.00 |
| barley (hordeum vulgare) | 4 | 24.67 | 0.99 | 48.00 |
| coarse barley kernel bread (80% scalded kernels, 20% white wheat flour) | 6 | 66.67 | 4.00 | 34.00 |
| beef (or other domesticated meat) | 10 | 0.00 | 0.00 | 70.00 |
| lentils and vegetables, eaten with orange | 15 | 10.50 | 1.58 | 35.00 |
| deer | 10 | 0.00 | 0.00 | 70.00 |
| Brown and Wild rice, Uncle Ben's Ready Whole Grain Medley (pouch) | 10 | 26.00 | 2.60 | 45.00 |
| chicken | 10 | 0.00 | 0.00 | 70.00 |
| raw vegetables salad (DE-1450) | 10 | 7.20 | 0.72 | 45.00 |
| TOTAL AGRICULTURE DIET: | | | 17.88 | 46.06 |

3000-0 YBP

| food item | % | g carb /100g serving | carbs contributed | GI |
|---|---|---|---|---|

| | | | | | | | |
|---|---|---|---|---|---|---|---|
| | | | | | | by food item | |
| | | | Moolgiri white jasmine rice (Tajmahal Agro Industries) | 20 | 21.33 | 4.27 | 54.00 |
| | | | steamed triticum aestivum | 5 | 74.63 | 3.73 | 55.00 |
| | | | barley (hordeum vulgare) | 4 | 24.67 | 0.99 | 48.00 |
| | | | coarse barley kernel bread (80% scalded kernels, 20% white wheat flour) | 6 | 66.67 | 4.00 | 34.00 |
| | | | raw vegetables salad (DE-1450) | 20 | 7.20 | 1.44 | 45.00 |
| | | | lentils and vegetables, eaten with orange | 15 | 10.50 | 1.58 | 35.00 |
| | | | chicken | 20 | 0.00 | 0.00 | 70.00 |
| | | | roasted bread from jowar flour (Sorghum vulgare) | 10 | 71.43 | 7.14 | 77.00 |
| | | | TOTAL AGRICULTURE DIET: | | | 23.14 | 55.69 |
| | | | TOTAL DIET, since 9000 YBP | | | 13.95 | 55.81 |

| | | | | | | | |
|---|---|---|---|---|---|---|---|
| | | | 9000-5000 YBP | | | | |
| UP Brahmins | North India | agriculture | food item | % in diet | g carb /100g serving | carbs contributed by food item | GI |
| | | | hunter-gatherer diet: | | | 6.35 | 60.45 |
| | | | 5000-4500 YBP | | | | |
| | | | food item | % | g carb /100g serving | carbs contributed by food item | GI |
| | | | (H-G diet) | 20 | 6.35 | 1.27 | 60.45 |
| | | | (H-G diet) | 5 | 6.35 | 0.32 | 60.45 |
| | | | (H-G diet) | 4 | 6.35 | 0.25 | 60.45 |
| | | | (H-G diet) | 6 | 6.35 | 0.38 | 60.45 |
| | | | (H-G diet) | 10 | 6.35 | 0.63 | 60.45 |
| | | | lentils and vegetables, eaten with orange | 15 | 10.50 | 1.58 | 35.00 |
| | | | deer | 10 | 0.00 | 0.00 | 70.00 |
| | | | Brown and Wild rice, Uncle Ben's Ready Whole Grain Medley (pouch) | 10 | 26.00 | 2.60 | 45.00 |

| | | | | |
|---|---|---|---|---|
| chicken | 10 | 0.00 | 0.00 | 70.00 |
| raw vegetables salad (DE-1450) | 10 | 7.20 | 0.72 | 45.00 |
| TOTAL AGRICULTURE DIET: | | | 7.75 | 48.66 |

4500-3000 YBP

| food item | % in diet | g carb /100g serving | carbs contributed by food item | GI |
|---|---|---|---|---|
| Moolgiri white jasmine rice (Tajmahal Agro Industries) | 20 | 21.33 | 4.27 | 54.00 |
| steamed triticum aestivum | 5 | 74.63 | 3.73 | 55.00 |
| barley (hordeum vulgare) | 4 | 24.67 | 0.99 | 48.00 |
| coarse barley kernel bread (80% scalded kernels, 20% white wheat flour) | 6 | 66.67 | 4.00 | 34.00 |
| beef (or other domesticated meat) | 10 | 0.00 | 0.00 | 70.00 |
| lentils and vegetables, eaten with orange | 15 | 10.50 | 1.58 | 35.00 |
| deer | 10 | 0.00 | 0.00 | 70.00 |
| Brown and Wild rice, Uncle Ben's Ready Whole Grain Medley (pouch) | 10 | 26.00 | 2.60 | 45.00 |
| chicken | 10 | 0.00 | 0.00 | 70.00 |
| raw vegetables salad (DE-1450) | 10 | 7.20 | 0.72 | 45.00 |
| TOTAL AGRICULTURE DIET: | | | 17.88 | 46.06 |

3000-0 YBP

| food item | % in diet | g carb /100g serving | carbs contributed by food item | GI |
|---|---|---|---|---|
| Moolgiri white jasmine rice (Tajmahal Agro Industries) | 25 | 21.33 | 5.33 | 54.00 |
| steamed triticum aestivum | 6 | 74.63 | 4.66 | 55.00 |
| barley (hordeum vulgare) | 5 | 24.67 | 1.23 | 48.00 |
| coarse barley kernel bread (80% scalded kernels, 20% white wheat flour) | 8 | 66.67 | 5.00 | 34.00 |

| | | | food item | % in diet | g carb /100g serving | carbs contributed by food item | GI |
|---|---|---|---|---|---|---|---|
| | | | beef (or other domesticated meat) | 13 | 0.00 | 0.00 | 70.00 |
| | | | lentils and vegetables, eaten with orange | 19 | 10.50 | 1.97 | 35.00 |
| | | | chicken | 13 | 0.00 | 0.00 | 70.00 |
| | | | raw vegetables salad (DE-1450) | 13 | 7.20 | 0.72 | 45.00 |
| | | | TOTAL AGRICULTURE DIET: | | | 18.92 | 46.25 |
| | | | TOTAL DIET, since 9000 YBP | | | 12.54 | 52.67 |

| | | | 9000-5000 YBP | | | | |
|---|---|---|---|---|---|---|---|
| North India tribe | North India | agriculture | food item | % in diet | g carb /100g serving | carbs contributed by food item | GI |
| | | | hunter-gatherer diet: | | | 6.35 | 60.45 |
| | | | 5000-4500 YBP | | | | |
| | | | food item | % | g carb /100g serving | carbs contributed by food item | GI |
| | | | (H-G diet) | 20 | 6.35 | 1.27 | 60.45 |
| | | | (H-G diet) | 5 | 6.35 | 0.32 | 60.45 |
| | | | (H-G diet) | 4 | 6.35 | 0.25 | 60.45 |
| | | | (H-G diet) | 6 | 6.35 | 0.38 | 60.45 |
| | | | (H-G diet) | 10 | 6.35 | 0.63 | 60.45 |
| | | | lentils and vegetables, eaten with orange | 15 | 10.50 | 1.58 | 35.00 |
| | | | deer | 10 | 0.00 | 0.00 | 70.00 |
| | | | Brown and Wild rice, Uncle Ben's Ready Whole Grain Medley (pouch) | 10 | 26.00 | 2.60 | 45.00 |
| | | | chicken | 10 | 0.00 | 0.00 | 70.00 |
| | | | raw vegetables salad (DE-1450) | 10 | 7.20 | 0.72 | 45.00 |
| | | | TOTAL AGRICULTURE DIET: | | | 7.75 | 48.66 |
| | | | 4500-0 YBP | | | | |
| | | | food item | % in diet | g carb /100g serving | carbs contributed by food item | GI |
| | | | Moolgiri white jasmine rice (Tajmahal Agro | 20 | 21.33 | 4.27 | 54.00 |

| | | | | | | | |
|---|---|---|---|---|---|---|---|
| | | | Industries) | | | | |
| | | | steamed triticum aestivum | 5 | 74.63 | 3.73 | 55.00 |
| | | | barley (hordeum vulgare) | 4 | 24.67 | 0.99 | 48.00 |
| | | | coarse barley kernel bread (80% scalded kernels, 20% white wheat flour) | 6 | 66.67 | 4.00 | 34.00 |
| | | | beef (or other domesticated meat) | 10 | 0.00 | 0.00 | 70.00 |
| | | | lentils and vegetables, eaten with orange | 15 | 10.50 | 1.58 | 35.00 |
| | | | deer | 10 | 0.00 | 0.00 | 70.00 |
| | | | Brown and Wild rice, Uncle Ben's Ready Whole Grain Medley (pouch) | 10 | 26.00 | 2.60 | 45.00 |
| | | | chicken | 10 | 0.00 | 0.00 | 70.00 |
| | | | raw vegetables salad (DE-1450) | 10 | 7.20 | 0.72 | 45.00 |
| | | | TOTAL AGRICULTURE DIET: | | | 17.88 | 46.06 |
| | | | TOTAL DIET, since 9000 YBP | | | 12.19 | 52.60 |

| | | | | | | | |
|---|---|---|---|---|---|---|---|
| | | | 9000-6000 YBP | | | | |
| Sindhi | Pakistan | agriculture | food item | % in diet | g carb /100g serving | carbs contributed by food item | GI |
| Hazara | | | lamb | 20 | 0.00 | 0.00 | 70.00 |
| Pathan | | | apricot raw (DE-135) | 15 | 7.50 | 1.13 | 57.00 |
| | | | steamed triticum aestivum | 12 | 74.63 | 8.96 | 55.00 |
| | | | barley (hordeum vulgare) | 8 | 24.67 | 1.97 | 48.00 |
| | | | dates, khalas (rutab, soft early ripened) variety | 15 | 50.00 | 7.50 | 47.00 |
| | | | beef | 20 | 0.00 | 0.00 | 70.00 |
| | | | Brown and Wild rice, Uncle Ben's Ready Whole Grain Medley (pouch) | 10 | 26.00 | 2.60 | 45.00 |
| | | | TOTAL PRE-6000 YBP DIET: | | | 22.15 | 22.15 |
| | | | 6000-4500 YBP | | | | |

| food item | % in diet | g carb /100g serving | carbs contributed by food item | GI |
|---|---|---|---|---|
| steamed triticum aestivum | 12 | 74.63 | 8.96 | 55.00 |
| barley (hordeum vulgare) | 9 | 24.67 | 2.22 | 48.00 |
| coarse barley kernel bread (80% scalded kernels, 20% white wheat flour) | 9 | 66.67 | 6.00 | 34.00 |
| lentils, type NS | 10 | 12.00 | 1.20 | 29.00 |
| TOTAL PRE-6000 YBP DIET: | 5 | 22.15 | 1.11 | 51.40 |
| TOTAL PRE-6000 YBP DIET: | 5 | 22.15 | 1.11 | 51.40 |
| TOTAL PRE-6000 YBP DIET: | 13 | 22.15 | 2.88 | 51.40 |
| lamb | 8 | 0.00 | 0.00 | 70.00 |
| apricot raw (DE-135) | 9 | 7.50 | 0.68 | 57.00 |
| dates, khalas (rutab, soft early ripened) variety | 10 | 50.00 | 5.00 | 47.00 |
| beef | 10 | 0.00 | 0.00 | 70.00 |
| TOTAL AGRICULTURE DIET: | | | 29.15 | 47.12 |

4500-0 YBP

| food item | % in diet | g carb /100g serving | carbs contributed by food item | GI |
|---|---|---|---|---|
| steamed triticum aestivum | 12 | 74.63 | 8.96 | 55.00 |
| barley (hordeum vulgare) | 9 | 24.67 | 2.22 | 48.00 |
| coarse barley kernel bread (80% scalded kernels, 20% white wheat flour) | 9 | 66.67 | 6.00 | 34.00 |
| lentils, type NS | 10 | 12.00 | 1.20 | 29.00 |
| Moolgiri white jasmine rice (Tajmahal Agro Industries) | 5 | 21.33 | 1.07 | 54.00 |
| chickpeas, boiled | 5 | 20.00 | 1.00 | 36.00 |
| whole milk (UK-277) | 13 | 4.50 | 0.59 | 34.00 |
| lamb | 8 | 0.00 | 0.00 | 70.00 |
| apricot raw (DE-135) | 9 | 7.50 | 0.68 | 57.00 |
| dates, khalas (rutab, soft early ripened) variety | 10 | 50.00 | 5.00 | 47.00 |

| | | | | | | | |
|---|---|---|---|---|---|---|---|
| | | | beef | 10 | 0.00 | 0.00 | 70.00 |
| | | | TOTAL AGRICULTURE DIET: | | | 26.70 | 45.87 |
| | | | TOTAL DIET, since 9000 YBP | | | 25.59 | 47.92 |

| | | | | | | | |
|---|---|---|---|---|---|---|---|
| | | | 9000-6000 YBP | | | | |
| Balochi | Pakistan | 0.66 agric, 0.33 pastoral | food item | % in diet | g carb /100g serving | carbs contributed by food item | GI |
| Brahui | | | (pre-6000 YBP diet) | | | 22.15 | 51.40 |
| Kalash | | | | | | | |
| Burusho | | | 6000-4500 YBP | | | | |
| | | | food item | % in diet | g carb /100g serving | carbs contributed by food item | GI |
| | | | (pre-6000 YBP diet) | 20 | 22.15 | 4.43 | 51.40 |
| | | | (pre-6000 YBP diet) | 5 | 22.15 | 1.11 | 51.40 |
| | | | lamb | 15 | 0.00 | 0.00 | 70.00 |
| | | | steamed triticum aestivum | 15 | 74.63 | 11.19 | 55.00 |
| | | | barley (hordeum vulgare) | 10 | 24.67 | 2.47 | 48.00 |
| | | | lentils, type NS | 5 | 12.00 | 0.60 | 29.00 |
| | | | apricot raw (DE-135) | 10 | 7.50 | 0.75 | 57.00 |
| | | | dates, khalas (rutab, soft early ripened) variety | 5 | 50.00 | 2.50 | 47.00 |
| | | | beef | 15 | 0.00 | 0.00 | 70.00 |
| | | | TOTAL PASTORAL DIET: | | | 23.05 | 51.91 |
| | | | 4500-0 YBP | | | | |
| | | | food item | % in diet | g carb /100g serving | carbs contributed by food item | GI |
| | | | whole milk (UK-277) | 20 | 4.50 | 0.90 | 34.00 |
| | | | chickpeas, boiled | 5 | 20.00 | 1.00 | 36.00 |
| | | | lamb | 15 | 0.00 | 0.00 | 70.00 |
| | | | steamed triticum aestivum | 15 | 74.63 | 11.19 | 55.00 |
| | | | barley (hordeum vulgare) | 10 | 24.67 | 2.47 | 48.00 |
| | | | lentils, type NS | 5 | 12.00 | 0.60 | 29.00 |
| | | | apricot raw (DE-135) | 10 | 7.50 | 0.75 | 57.00 |

| | | | | | | | |
|---|---|---|---|---|---|---|---|
| | | | dates, khalas (rutab, soft early ripened) variety | 5 | 50.00 | 2.50 | 47.00 |
| | | | beef | 15 | 0.00 | 0.00 | 70.00 |
| | | | TOTAL PASTORAL DIET: | | | 19.41 | 50.40 |
| | | | PASTORAL, since 9000 YBP | | | 20.93 | 50.99 |
| | | | AGRICULTURE, since 9000 YBP | | | 25.59 | 47.92 |
| | | | TOTAL DIET, since 9000 YBP | | | 24.04 | 48.94 |

| | | | | | | | |
|---|---|---|---|---|---|---|---|
| | | | 9000-6000 YBP | | | | |
| Makrani | Pakistan | cultivator | food item | % in diet | g carb /100g serving | carbs contributed by food item | GI |
| | | | (pre-6000 YBP diet) | | | 22.15 | 51.40 |
| | | | 6000-4500 YBP | | | | |
| | | | food item | % in diet | g carb /100g serving | % carbs contributed by food item | GI |
| | | | steamed triticum aestivum | 15 | 74.63 | 11.19 | 55.00 |
| | | | barley (hordeum vulgare) | 5 | 24.67 | 1.23 | 48.00 |
| | | | (pre-6000 YBP diet) | 8 | 22.15 | 1.77 | 51.40 |
| | | | (pre-6000 YBP diet) | 5 | 22.15 | 1.11 | 51.40 |
| | | | lentils, type NS | 10 | 12.00 | 1.20 | 29.00 |
| | | | apricot raw (DE-135) | 14 | 7.50 | 1.05 | 57.00 |
| | | | dates, khalas (rutab, soft early ripened) variety | 13 | 50.00 | 6.50 | 47.00 |
| | | | (pre-6000 YBP diet) | 14 | 22.15 | 3.13 | 51.40 |
| | | | lamb | 8 | 0.00 | 0.00 | 70.00 |
| | | | beef | 8 | 0.00 | 0.00 | 70.00 |
| | | | 6000-4500 YBP diet: | | | 27.19 | 50.90 |
| | | | 4500-3000 YBP | | | | |
| | | | food item | % in diet | g carb /100g serving | carbs contributed by food item | GI |

| | | | | |
|---|---|---|---|---|
| steamed triticum aestivum | 15 | 74.63 | 11.19 | 55.00 |
| barley (hordeum vulgare) | 5 | 24.67 | 1.23 | 48.00 |
| 6000-4500 YBP diet: | 8 | 22.15 | 1.77 | 51.40 |
| Moolgiri white jasmine rice (Tajmahal Agro Industries) | 5 | 21.33 | 1.07 | 54.00 |
| lentils, type NS | 10 | 12.00 | 1.20 | 29.00 |
| apricot raw (DE-135) | 14 | 7.50 | 1.05 | 57.00 |
| dates, khalas (rutab, soft early ripened) variety | 13 | 50.00 | 6.50 | 47.00 |
| whole milk (UK-277) | 14 | 4.50 | 0.63 | 34.00 |
| lamb | 8 | 0.00 | 0.00 | 70.00 |
| beef | 8 | 0.00 | 0.00 | 70.00 |
| TOTAL CULTIVATOR DIET: | | | 24.65 | 50.52 |

3000-0 YBP

| food item | % in diet | g carb /100g serving | carbs contributed by food item | GI |
|---|---|---|---|---|
| steamed triticum aestivum | 15 | 74.63 | 11.19 | 55.00 |
| barley (hordeum vulgare) | 5 | 24.67 | 1.23 | 48.00 |
| roasted bread from jowar flour (Sorghum vulgare) | 8 | 71.43 | 5.71 | 77.00 |
| Moolgiri white jasmine rice (Tajmahal Agro Industries) | 5 | 21.33 | 1.07 | 54.00 |
| lentils, type NS | 10 | 12.00 | 1.20 | 29.00 |
| apricot raw (DE-135) | 14 | 7.50 | 1.05 | 57.00 |
| dates, khalas (rutab, soft early ripened) variety | 13 | 50.00 | 6.50 | 47.00 |
| whole milk (UK-277) | 14 | 4.50 | 0.63 | 34.00 |
| lamb | 8 | 0.00 | 0.00 | 70.00 |
| beef | 8 | 0.00 | 0.00 | 70.00 |
| TOTAL CULTIVATOR DIET: | | | 28.59 | 55.76 |
| TOTAL CULTIVATOR, since 9000 YBP | | | 25.55 | 52.62 |

**Supplementary Table 10.** List of populations included in the analysis, their locations, traditional occupations and estimation of average diet over the last 9000 years as measured by GI and % carbohydrates, as constructed in Diet model 2. Summary of Supplementary Table 9. Ancestral group refers to the assigned population grouping for BayeScan and BayeScEnv analyses, defined by ADMIXTURE.

| Ancestral group | Population | Location | N | Latitude | Longitude | Occupation | GI | Carbohydrate content |
|---|---|---|---|---|---|---|---|---|
| 1 | Brahmins_UP | India | 56 | 26.06 | 83.18 | Agriculture | 52.67 | 12.54 |
| 1 | North India caste | India | 54 | 24.39 | 82.45 | Agriculture | 52.67 | 12.54 |
| 1 | North India tribe | India | 19 | 27.87 | 78.98 | Agriculture | 52.6 | 12.19 |
| 1 | Central India tribe | India | 7 | 21.3 | 80.46 | Hunter-gatherer | 60.45 | 6.35 |
| 1 | Gujarati | India | 88 | 23.32 | 72.31 | Agriculture | 55.81 | 13.95 |
| 1 | South India caste | India | 30 | 15.35 | 77.57 | Agriculture | 54.98 | 17.19 |
| 1 | South India tribe | India | 24 | 13.19 | 76.7 | Hunter-gatherer | 60.45 | 6.35 |
| 2 | Brahui | Pakistan | 25 | 30.5 | 66.5 | 2/3 agric, 1/3 pastoral | 48.94 | 24.04 |
| 2 | Balochi | Pakistan | 25 | 30.5 | 66.5 | 2/3 agric, 1/3 pastoral | 48.94 | 24.04 |
| 2 | Makrani | Pakistan | 25 | 26 | 64 | Cultivator | 52.62 | 25.55 |
| 2 | Sindhi | Pakistan | 25 | 25.5 | 69 | Agriculture | 47.92 | 25.59 |
| 2 | Pathan | Pakistan | 23 | 33.5 | 70.5 | Agriculture | 47.92 | 25.59 |
| 2 | Kalash | Pakistan | 25 | 36 | 70.5 | 2/3 agric, 1/3 pastoral | 48.94 | 24.04 |
| 2 | Burusho | Pakistan | 25 | 36.5 | 74 | 2/3 agric, 1/3 pastoral | 48.94 | 24.04 |
| 3 | Han | China | 44 | 32.5 | 114 | Agriculture | 62.15 | 14.76 |
| 3 | Miaozu | China | 10 | 28 | 109 | Agriculture | 62.15 | 14.76 |
| 3 | Naxi | China | 9 | 26 | 100 | 2/3 agric, 1/3 pastoral | 58.43 | 10.37 |
| 3 | She | China | 10 | 27 | 119 | Agriculture | 62.15 | 14.76 |
| 3 | Tu | China | 10 | 36 | 101 | Pastoral | 56.82 | 8.65 |
| 3 | Tujia | China | 10 | 29 | 109 | Agriculture | 62.15 | 14.76 |
| 3 | Xibo | China | 9 | 43.5 | 81.5 | 1/3 agric, 2/3 pastoral | 45.62 | 7.75 |

| | | | | | | | | |
|---|---|---|---|---|---|---|---|---|
| 3 | Yizu | China | 10 | 28 | 103 | 1/3 agric, 2/3 pastoral | 58.07 | 10.64 |
| 3 | East India Sino-Tibetan speakers | India | 8 | 26.08 | 92.48 | Hunter-gatherer | 60.45 | 6.35 |
| 3 | Japanese | Japan | 30 | 38.1 | 138.3 | Agriculture | 55.48 | 10.96 |
| 3 | Cambodian | Cambodia | 11 | 12 | 105 | Agriculture | 59.31 | 12.98 |
| 3 | Burmese | Myanmar | 15 | 23.52 | 95.75 | Agriculture | 55.95 | 9.79 |
| 3 | Dai | Myanmar | 10 | 21 | 100 | Agriculture | 56.7 | 12.03 |
| 4 | Hazara | Pakistan | 24 | 33.5 | 70 | Agriculture | 47.92 | 25.59 |
| 4 | Uygur | China | 10 | 44 | 81 | Agriculture | 46.37 | 12.29 |

**Supplementary Table 11**. Populations included in the South Asian population groupings

| Population group | Population | Source | *N* | Latitude | Longitude |
|---|---|---|---|---|---|
| **Central_tribe** | | | **7** | **21.3** | **80.46** |
| | Gond | Metspalu et al. (2011) | 3 | 22.1 | 82.14 |
| | Bhunjia | Metspalu et al. (2011) | 1 | 18.5 | 82.5 |
| | Nihali | Metspalu et al. (2011) | 1 | 21.23 | 81.63 |
| | Dhurwa | Metspalu et al. (2011) | 2 | 21.54 | 76.33 |
| **EastIN_AA** | | | **23** | **21.88** | **84.9** |
| | Asur | G. Chaubey | 2 | 24 | 84.1 |
| | Bonda | G. Chaubey | 4 | 18.5 | 82.5 |
| | Gadaba | G. Chaubey | 1 | 18.5 | 82.5 |
| | Ho | G. Chaubey | 5 | 25.42 | 86.13 |
| | Juang | G. Chaubey | 2 | 21.28 | 84.4 |
| | Kharia | G. Chaubey | 2 | 21.9 | 83.4 |
| | Khasi | G. Chaubey | 3 | 25.5 | 90.33 |
| | Mawasi | G. Chaubey | 1 | 12 | 84.7 |
| | Santhal | G. Chaubey | 1 | 23.08 | 84.66 |
| | Savara | G. Chaubey | 2 | 18.82 | 82.72 |
| **EastIN_ST** | | | **12** | **26.08** | **92.48** |
| | Garo | G. Chaubey | 4 | 25.3 | 90.3 |
| | Naga | G. Chaubey | 4 | 25.67 | 94.11 |
| | Nyishi | G. Chaubey | 4 | 27.26 | 93.03 |
| **NorthIN_tribe** | | | **19** | **27.87** | **78.98** |
| | Kol | Metspalu et al. (2011) | 16 | 27.9 | 78.13 |
| | Tharus | G. Chaubey | 3 | 27.7 | 83.46 |

| | | | | | |
|---|---|---|---|---|---|
| **NorthIN_caste** | | | **54** | **24.39** | **82.45** |
| | Chamar | Metspalu et al. (2011) | 10 | 25.28 | 82.95 |
| | Dharkars | Metspalu et al. (2011) | 12 | 25.28 | 82.95 |
| | Dusadh | Metspalu et al. (2011) | 10 | 18 | 85.01 |
| | Kanjars | Metspalu et al. (2011) | 9 | 26.47 | 80.35 |
| | Kshatriya | Metspalu et al. (2011) | 7 | 27.63 | 78.67 |
| | Kurmi | Metspalu et al. (2011) | 1 | 22.57 | 88.36 |
| | UP_lowcaste | G. Chaubey | 5 | 25.37 | 83.04 |
| **Brahmins_UP** | | G. Chaubey | **56** | **26.06** | **83.18** |
| **SouthIN_tribe** | | | **24** | **13.19** | **76.7** |
| | Chenchus | G. Chaubey | 4 | 17.26 | 80.09 |
| | Halakipikki | G. Chaubey | 5 | 14.6 | 74.7 |
| | Kurumba | Metspalu et al. (2011) | 4 | 11.4 | 76.88 |
| | Lambadi | Metspalu et al. (2011) | 1 | 17.37 | 78.48 |
| | Malayan | Behar et al. (2010) | 2 | 11.51 | 76.02 |
| | Paniya | Behar et al. (2010) | 4 | 11.64 | 76.01 |
| | Sakilli | Behar et al. (2010) | 4 | 10.52 | 76.21 |
| **SouthIN_caste** | | | **30** | **15.35** | **77.57** |
| | N. Kannadi | Behar et al. (2010) | 9 | 15.46 | 75.01 |
| | Pulliyar | Metspalu et al. (2011) | 5 | 11.02 | 76.97 |
| | Brahmins_AP | Metspalu et al. (2011) | 4 | 17.39 | 78.49 |
| | Velmas | G. Chaubey | 10 | 17.05 | 79.27 |
| | TN_Low Caste | Metspalu et al. (2011) | 2 | 13.06 | 80.25 |
| **Gujaratis** | | HapMap 3 | **88** | **23.31** | **72.31** |

**Supplementary Table 12.** Top ontology terms that overlapped among several variables and across methods. Because GREAT incorporates gene regions, many of the genes in the ontology term may not have SNPs represented in the significant regions of interest. Therefore, we only highlighted genes which contained statistically significant SNPs. In instances where there were multiple genes to highlight, we prioritized those containing SNPs with published gene or allele-specific expression data. Lastly some of these genes appear to be associated with other diet variables - however, we only report conditions under which these genes appear in conjunction with the associated ontology term. In parentheses beside the SNP IDs are the specific variables associated with each SNP: a refers to vitamin A, p to protein, l to lipids, z to zinc. Baypass means that the SNP was only associated with Baypass results, not BayeScEnv or BayeScan.

| **Dietary variable** | **Ontology term** | **Methods** | **Gene** | **SNP** |
|---|---|---|---|---|
| Vitamin A | Loop of Henle development | All three tests | N/A | No specified SNP |
| Vitamin A | Negative regulation of type B pancreatic cell apoptotic process | BayeScan, BayeScEnv | *CAST* | rs152019 |
| | | | *TCF7L2* | rs10885409 |

| | | | | |
|---|---|---|---|---|
| Omega 3 | Positive regulation of type B pancreatic cell apoptotic process | Baypass | *IL6* | rs2066992 |
| GI | | | *N/A* | No specified SNP |
| Vitamin A | Response to food | BayeScEnv | *GHR* | rs4292454 (a) |
| Omega 3 | | Baypass | | rs4129472, rs4292454 |
| GL | | | | rs13153388, rs4242116, rs7701605 |

| | | | | |
|---|---|---|---|---|
| Lipids | Branching involved in salivary gland morphogenesis | BayeScEnv | *LAMA1* | rs4798530 (p,l) rs11873300 (p, l, z) rs16951201 (p, l, z) |
| Protein | | | | |
| Zinc | | | *SEMA3A* | rs17158741 (p,l,z) Rs16887725 (p,l, z) |
| Omega 6 | | BayeScEnv/ Baypass | *LAMA1* | rs4798530, rs11873300, rs16951201, rs633691 (Baypass) |
| | | | *SEMA3A* | rs16887725 rs17158741 |
| GI | | Baypass | *LAMA1* | rs648161 |
| Omega 3 | | Baypass | *LAMA1* | rs4239330 |

**Supplementary Figure 1. Distribution of HGDP populations across Asia.** a) ADMIXTURE results and geographic location of populations. ADMIXTURE results show ancestry groups with specified *K*=3 or *K*=4 groupings. The numbers above the *K*=3 plot refer to the population grouping used for BayeScan and BayeScEnv analyses, and the colors in the map reflect these groupings**.** b) The colors of the large triangles refer to population subsistence strategy. In purple are the known locations of rice domestication, through time.

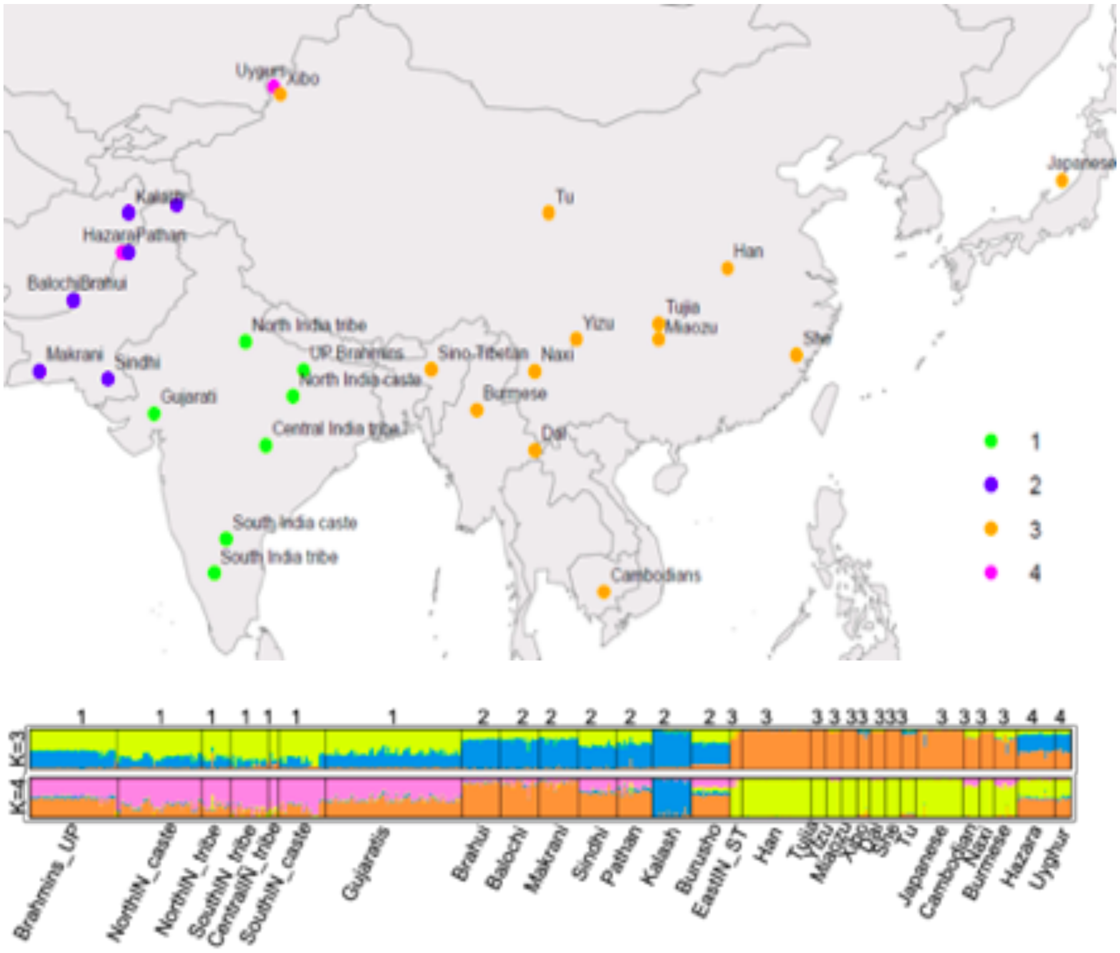


Supplementary Figure 1a.

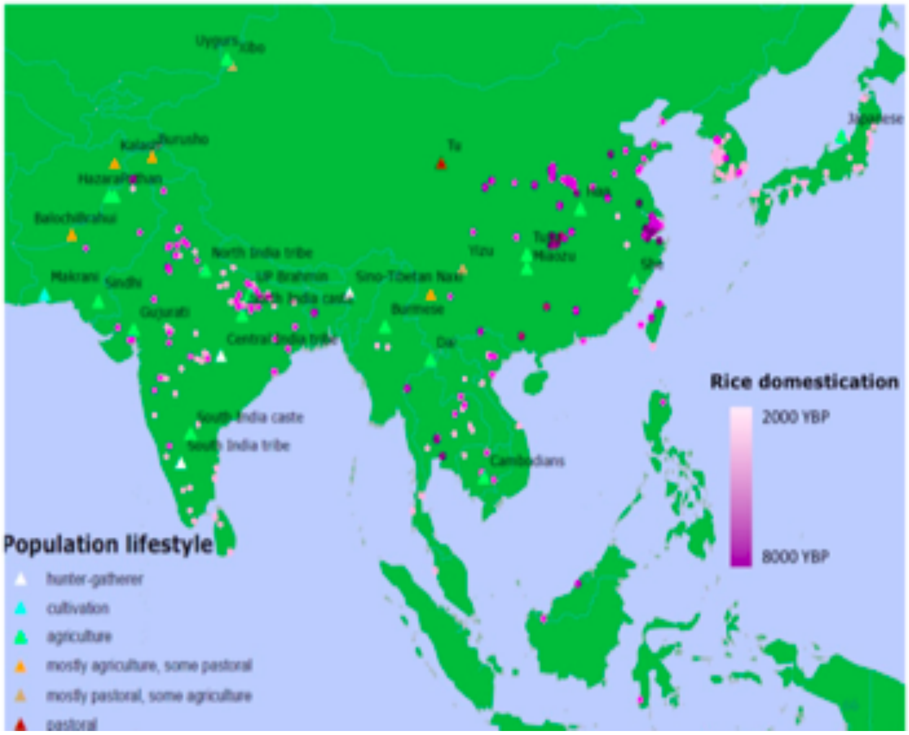


Supplementary Figure 1b.

**Supplementary Figure 2. Overlap between selection results.** Venn diagrams of the overlaps between SNPs in the top 1% (Baypass, BayeScan) or 0.01 *q*-value (BayeScEnv) of gene-environment correlations for each diet variable. Iron did not have any statistically significant hits for BayeScEnv, so only overlap between Baypass and BayeScan results are provided.

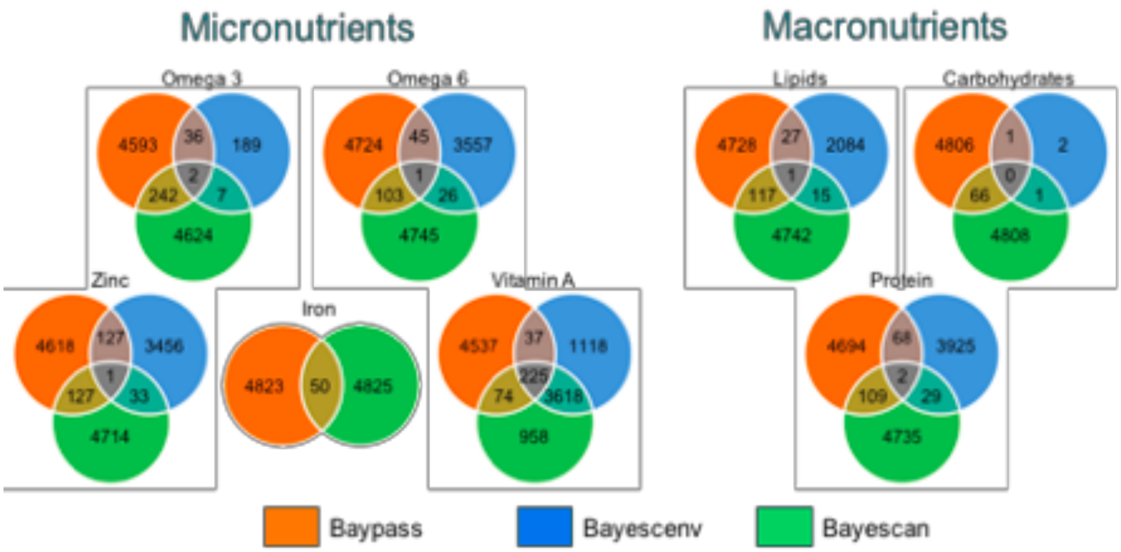


**Supplementary Figure 3.** Identification of known genes containing variants that are reported to be under positive selection in human populations, in the top 1% of results for Baypass, BayeScEnv and BayeScan. Note that as BayeScan does not incorporate environmental information, all variables are highlighted as showing association with the gene of interest in BayeScan results.

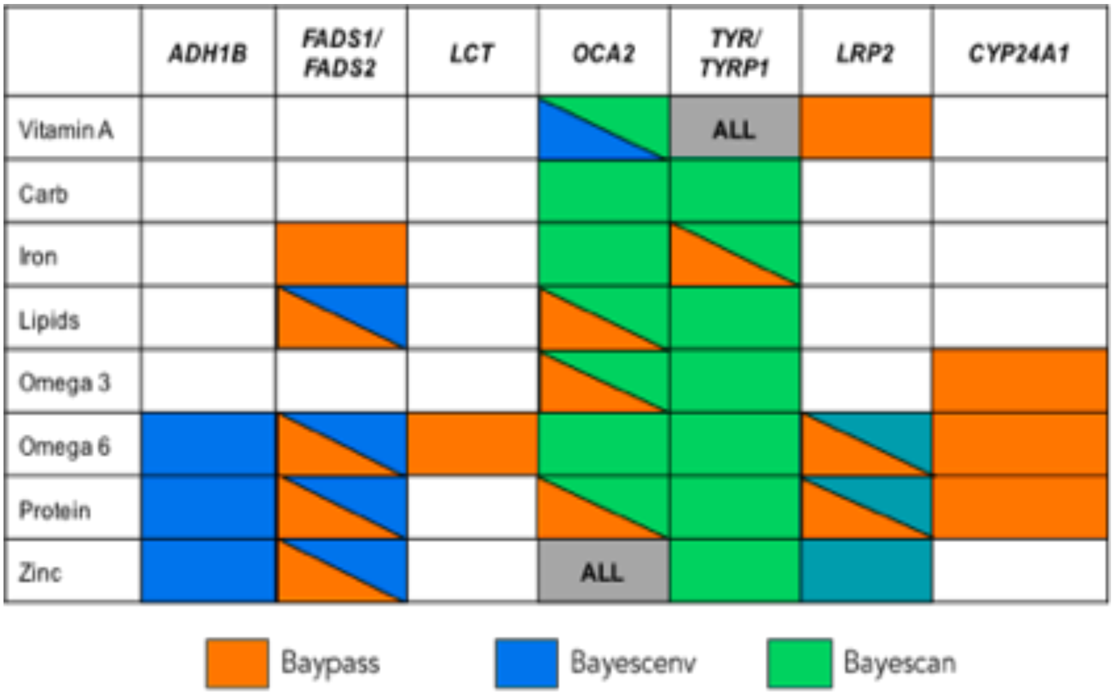


**Supplementary Figure 4. Annotated functions of top 1% of selection results, per nutrient and selection method.** On the y-axis are percentages of SNPs in each category.

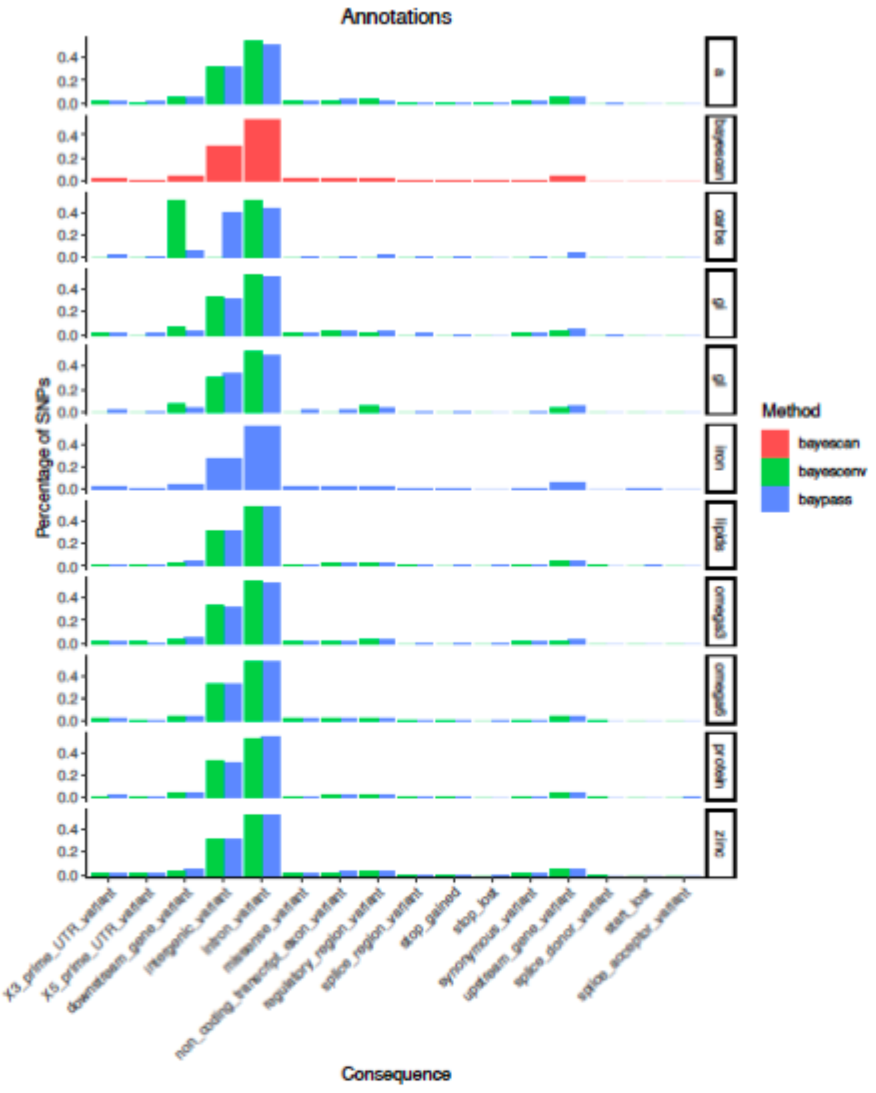


**Supplementary Figure 5. Numbers of SNPs classified as deleterious based on PolyPhen and SIFT scores.**

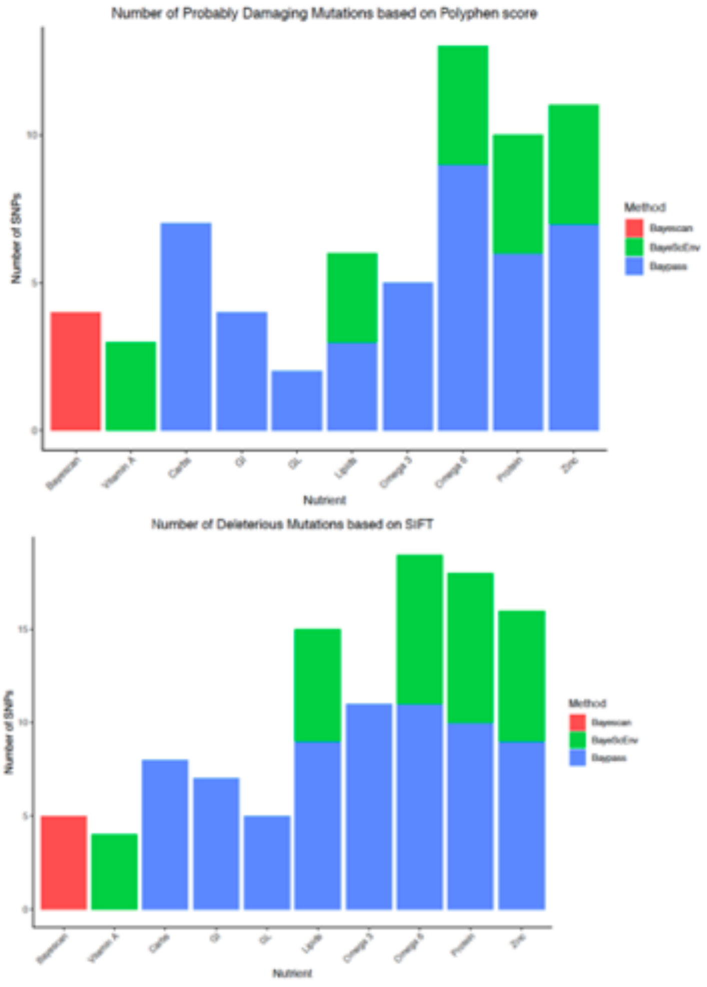


**Supplementary Figure 6. Enrichment of GWAS results in all three selection tests, for each nutrient variable.** Each panel depicts the enrichment results for each scan, with size of the circle corresponding to the combined “Odds ratio” of the SNPs under selection associated with each trait. The color of each point refers to the q-value for significance of the enrichment, based on a binomial test, the entire genome (1000 genomes, CEU) as background, and 30,553 SNPs associated with 573 traits downloaded November 1, 2018 from the NIH GWAS database.

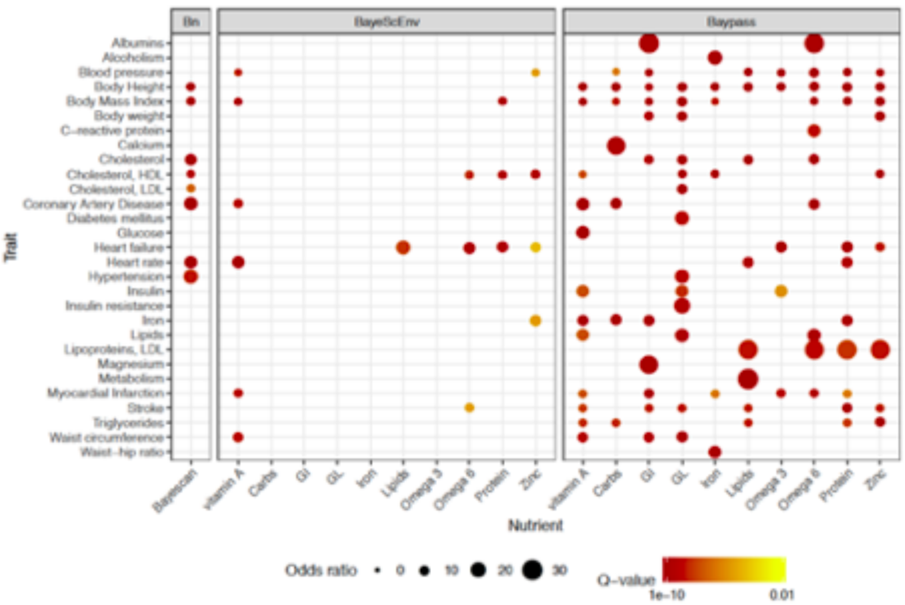


**Supplementary Figure 7.** Expression as measured by FPKM values (Fragments Per Kilobase of transcript per Million mapped reads), from Martin et al. 2014). Each dot represents the FPKM values of a single individual from the specified population.

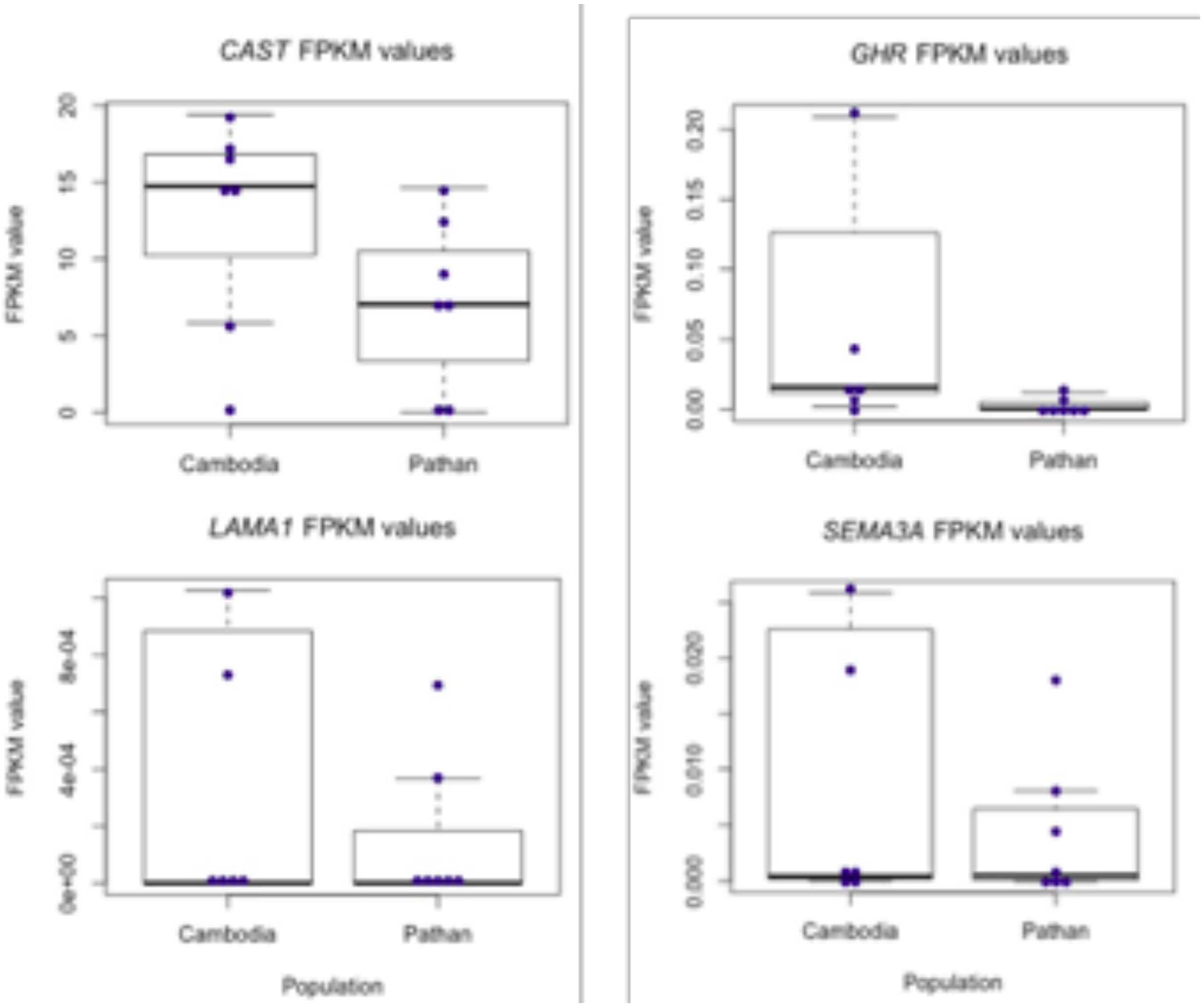

CAST FPKM values
GHR FPKM values
LAMA1 FPKM values
SEMA3A FPKM values
FPKM value
Cambodia
Pathan
Population

**Supplementary Figure 8.** Allele-specific expression differences in SNPs located in the CAST and GHR genes. The values are based on the difference between weighted reference allele counts and unweighted reference allele counts. The All column refers to the entire sample used in Martin et al. (2014), whereas Pathan and Cambodia groups represent subsections of that larger dataset.

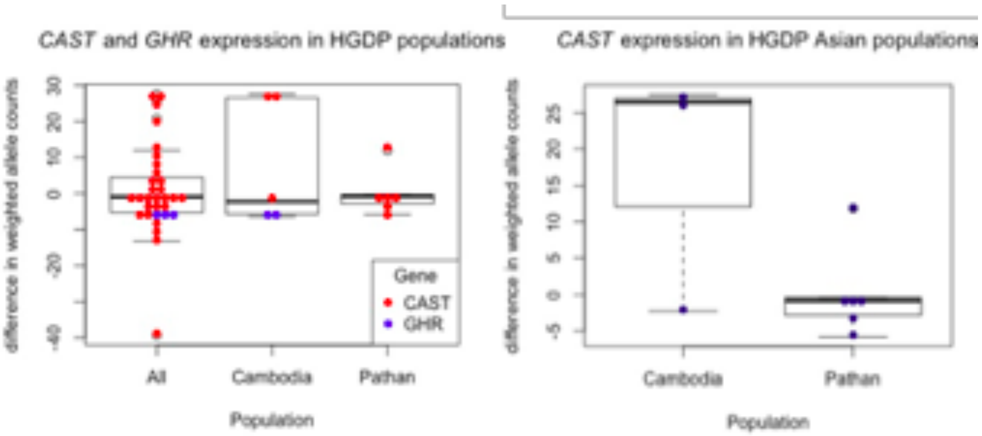

**Supplementary Figure 9.** SNP derived allele frequencies in Asian populations, against interpolated values of correlated dietary variables from Diet Model 1 - for the *LAMA1* gene, against 4 dietary variables.

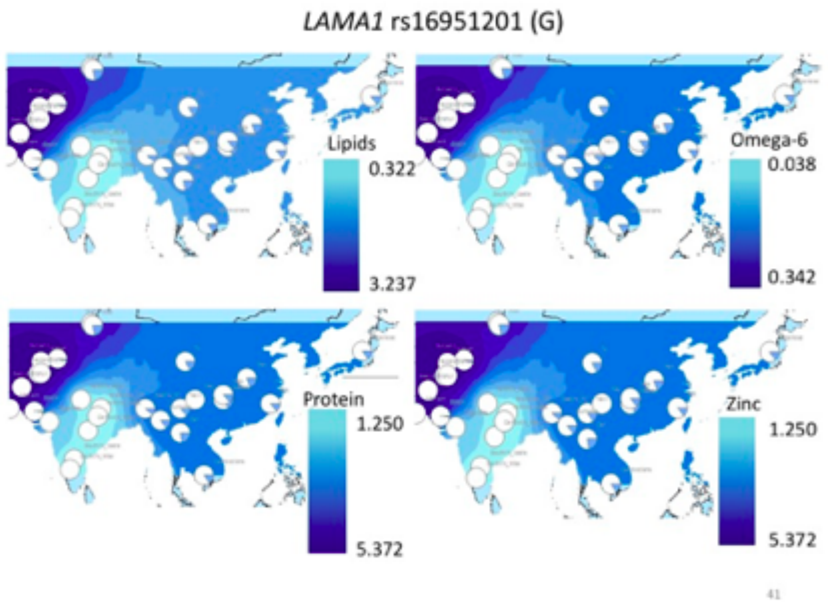

**Supplementary Figure 10.** SNP derived allele frequencies in Asian populations, against interpolated values of correlated dietary variables from Diet Model 1 - for the *SEMA3A* gene, against 4 dietary variables.

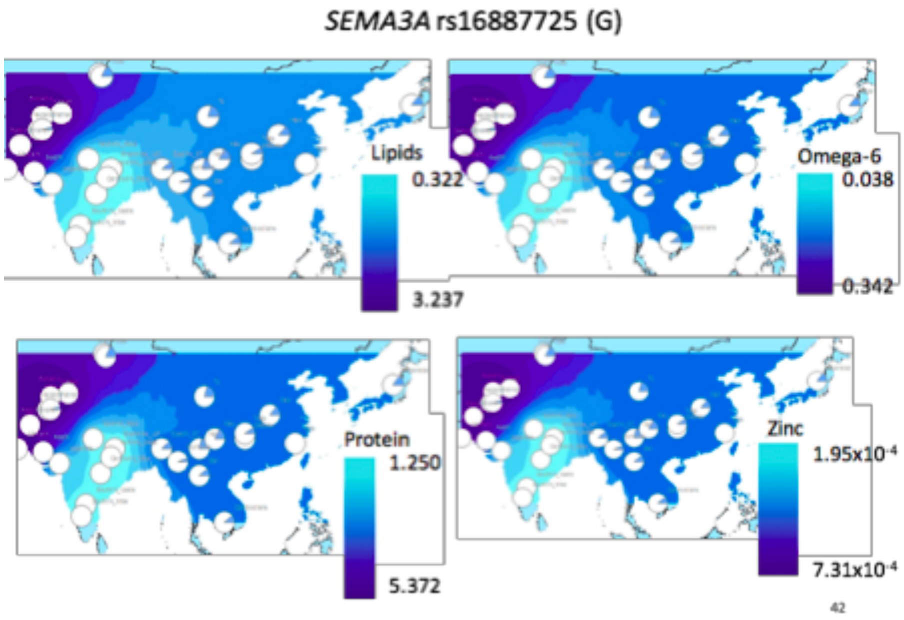
